# Distinct neurogenic pathways shape the diversification and mosaic organization of cortical output channels

**DOI:** 10.1101/2025.07.18.665624

**Authors:** Shreyas M. Suryanarayana, Xu An, Yongjun Qian, Shengli Zhao, Hemanth Mohan, Z. Josh Huang

## Abstract

The cerebral cortex broadcasts its output to subcortical regions through the projections of diverse extratelencephalic neurons (ETs) derived from either direct (dNG-ET^d^) or indirect (iNG-ET^i^) neurogenesis, but the differential contributions of these neurogenic pathways to cortical output organization remain unknown. Establishing a lineage-based viral targeting strategy in mice, we uncover that ET^i^ massively amplifies and diversifies ET^d^ projections across the cortex. Whereas ET^d^ primarily targets the forebrain and midbrain, ET^i^ dominates the hindbrain (e.g., brainstem/spinal action centers, lemniscal sensory stations), hypothalamic, and neuromodulatory regions with cortical area-specific projections. Numerous corticofugal subpopulations derive solely from iNG-ET^i^. Area-specific spinal projections of ET^i^ but not ET^d^ emerge from the postnatal pruning of an initial cortex-wide population. In the motor cortex, ET^d^ or ET^i^ activation induced distinct movements: either head-trunk, or additionally, coordinated orofacial-forelimb movements, respectively. Thus, two foundational neurogenic pathways with distinct evolutionary history differentially shape the area-specific diversification of cortical output channels.

## Introduction

The cerebral cortex is the crowning achievement of vertebrate evolution and contains a vast network of nerve cells underlying high-level brain functions from sensory processing to cognition and motor control ^1–4^. Glutamatergic projection neurons (PN) constitute up to 80% of cortical neurons and comprise several major classes characterized by their laminar location, projection patterns, and gene expression profiles ^5–7^. Whereas the intratelencephalic (IT) neurons project within the cerebral hemisphere and mediate intra-cortical processing streams, the extratelencephalic (ET) neurons constitute a massive and area specific efferent system that broadcast cortical output to myriad subcortical regions throughout the central nervous system ^6,8–10^. Decades of studies have uncovered daunting diversity in ET projection patterns within and across cortical areas and across mammalian species ^10–15^. Nevertheless, the same diverse set of ET neurons are reliably generated in each animal to carry out species-typic behavioral functions, suggesting a robust and stringent developmental genetic program. However, the development origins and processes that shape ET projection diversity and their area specific profiles remain poorly understood.

All PNs, including ETs, are generated from radial glia progenitors (RG) lining the embryonic cerebral ventricle through two fundamental forms of neurogenesis. In direct neurogenesis (dNG), a RG undergoes asymmetric division to self-renew and generate one neuronal progeny^4,16,17^; in indirect neurogenesis (iNG), RG first produces an intermediate progenitor (IP), which then undergoes symmetric division to generate two neurons^18–24^. Across the developing mammalian neural tube, dNG is ubiquitous from spinal segments to the forebrain and contributes to the development of the entire central nervous system, while iNG is largely restricted to the telencephalon that gives rise to the cerebral hemisphere^18,25^. Across vertebrate evolution, while dNG originated before the dawn of vertebrates and has been conserved ever since, iNG emerged largely since amniotes (but see ^26,27^) and has expanded tremendously in mammals, and is thought to drive the evolutionary innovation of a 6-layered isocortex^20,23,25,28,29^. While the amplification of neuronal production through IP is inherent to iNG, whether and how dNG and iNG differentially contribute to the diversification and organization of ET-mediated cortical output channels is unclear.

Using a genetic strategy to distinguish and fate map dNG and iNG, we previously found that whereas dNG generates all major PN classes, iNG differentially amplifies and diversifies PNs within each class ^17^. In particular, dNG and iNG generate ET^d^ and ET^i^, respectively, which are completely intermixed in layers 5 and 6 and appear indistinguishable in gross somato-dendritic morphology^17,30^. However, whether ET^d^ and ET^i^ differ in axon projection patterns - a crucial determinant of their connectivity and circuit function, remains unknown.

Here, we first establish a novel lineage-based genetic method that enables differential viral targeting of ET^d^ and ET^i^ in the same individual mice. Through systematic antero- and retrograde tracing, we show that iNG-ET^i^ massively amplifies and diversifies the dNG-ET^d^ projections across the cortex. While ET^d^ projections are largely restricted to forebrain and midbrain structures, ET^i^ neurons greatly amplify and diversify these projections and overwhelmingly dominate the innervation of hindbrain and spinal cord in an area-specific pattern. In particular, ETs^i^ in motor areas dominate projections to brainstem and spinal action diversification and execution centers, in sensory areas expand projections to sensory processing stations along lemniscal pathways, in higher order areas extend projections to other pallial structures and to hypothalamic and neuromodulatory centers. Corticofugal subpopulations in multiple areas are derived from only ET^i^ but not ET^d^, indicating the generation of novel projection types by iNG. During an early postnatal period, ET^i^ but not ET^d^ neurons across the cortex undergo exuberant axon overgrowth to the spinal cord followed by massive pruning, sculpting areal-specific spinal projections seen in mature cortex. Optogenetic activation reveals functional distinction between ET^d^ and ET^i^ in the motor cortex: while the formal induced movements of head and trunk, the latter additionally elicited coordinated orofacial and forelimb movements that resemble feeding. Together, these results demonstrate that two foundational neurogenic pathways with distinct evolutionary history prime the area-specific diversification and mosaic organization of cortical output channels that shape the central nervous system architecture.

## Results

### A genetic strategy enabling differential viral access to ET^d^ and ET^i^ in the same animal

To fully uncover the anatomical distinctions between ET^d^ and ET^i^, we designed a novel genetic strategy that allows for differential viral targeting of these two subpopulations based on their lineage distinction that resolves a common molecular marker (**Figure 1A**). The zinc finger transcription factor *Fezf2* is broadly expressed in the majority of postmitotic ETs including both ET^d^ and ET^i^ ^17,31,32^, yet the T-box transcription factor *Tbr2* is selectively expressed in embryonic IPs that generate all PN^i^, including ET^i^ but not ET^d^ ^17^. Thus, we built a *FezF2-tTA-CreER (FezF2-TC)* knockin allele driving the conditional expression of two tandemly arranged cassettes: a tTA cassette flanked by Frt sequences followed by a CreER cassette (**Figure 1A**). We confirmed that *FezF2-TC* reliably captures endogenous *FezF2* expression pattern by *in situ* detection of *FezF2* mRNA in labelled ET neurons (**Figure S1A-C**). When combined with a *Tbr2-Flp* knockin allele ^17^, dNG-derived ETs^d^, which do not express *Tbr2-Flp* during development, constitutively express tTA but not CreER; on the other hand, iNG and IP-derived ETs^i^, in which developmental expression of *Tbr2-Flp* removes the tTA cassette, constitutively express CreER (**Figure 1A**). Thus, this genetic strategy converts developmentally transient lineage signals in neurogenesis to permanent “genetic handles” that allow differential viral targeting of postmitotic ET^d^ and ET^i^ neurons in the same individual mice.

**Figure 1.**
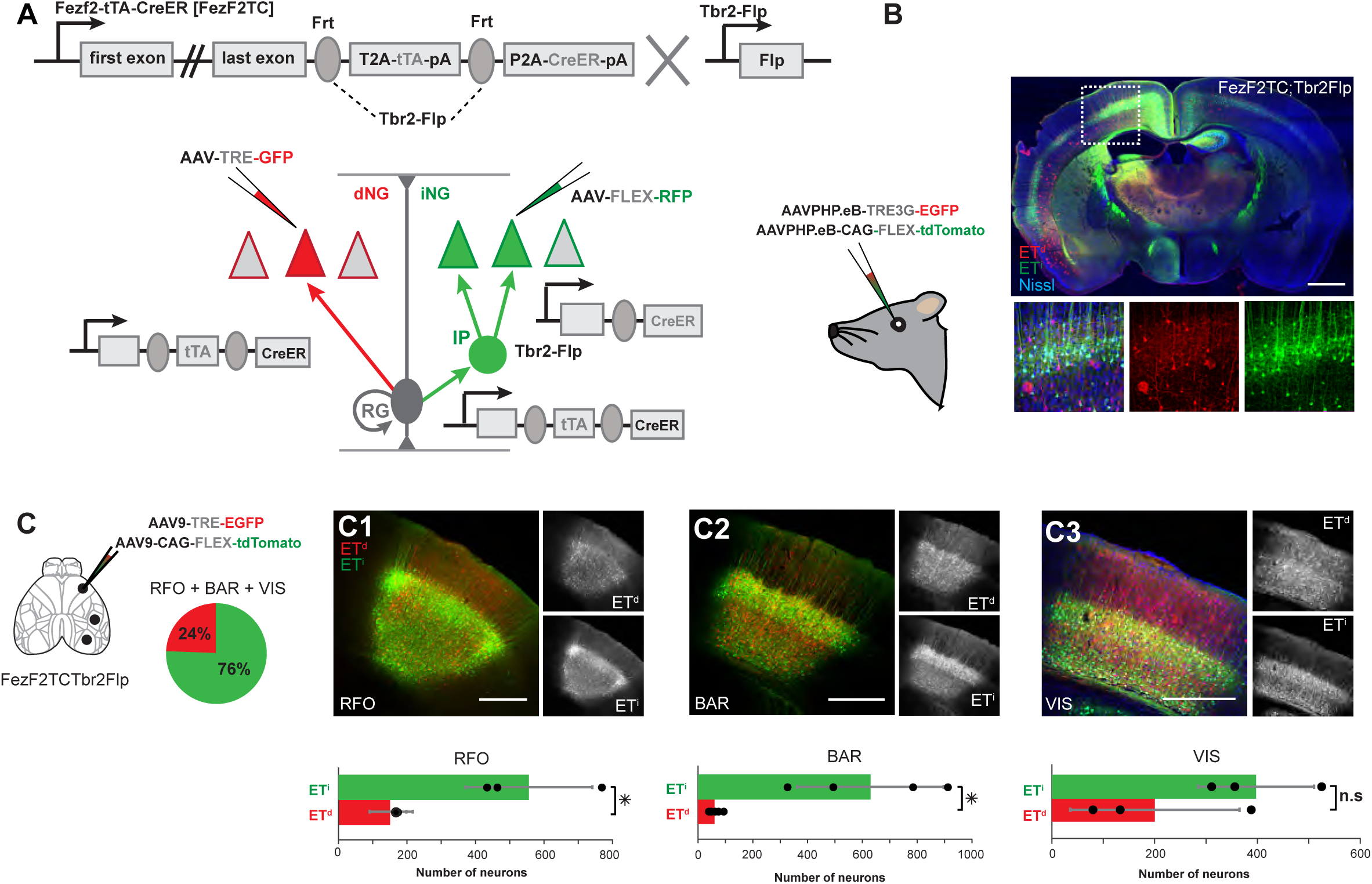
Genetic strategy for differential targeting of *Fezf2^+^* ET^d^ and ET^i^ PNs in the same mouse. (A) Schematic of *Fezf2-tTA-CreER (FezF2TC)* knockin allele where a *frt-tTA-frt-CreER* cassette was inserted in frame before the STOP codon of *Fezf2* gene. In *FezfTC;Tbr2-Flp* double*-*allele mice, dNG-derived ET^d^ express *tTA* while iNG-derived ET^i^ express *CreER*, as *tTA* is removed by *Tbr2-Flp* in IPs. Postnatal injection of *AAV-TRE-GFP* and *AAV-FLEX-RFP* in cortex differentially label ET^d^ and ET^i^, respectively. (B) Cortex wide labelling of ET^d^ and ET^i^ neurons through retro-orbital co-injections of *AAVPHP.eB-TRE-EGFP* and *AAVPHP.eB-FLEX-tdTomato* in *FezfTC;Tbr2-Flp* mice. (C) Schematic of stereotactic AAV injection to label ET^d^ and ET^i^ using *FezF2-TC;Tbr2Flp* mice in the RFO, BAR, and VIS. Labeled ET neurons across the three areas consist of 24% ET^d^ and 76% ET^i^. (C1-C3) Focal labelling of ET^d^ (red) and ET^i^ (green) in RFO (C1) BAR (C2), and VIS (C3). Significantly more number of ET^i^ than ET^d^ neurons were labelled in all three areas (bottom panels). Abbreviations: RFO – rostral forelimb orofacial area in motor cortex; BAR – barrel cortex; VIS – visual cortex; dNG – direct neurogenesis; iNG - indirect neurogenesis; ET^d^ – direct neurogenesis generated extratelencephalic neuron; ET^i^ – indirect neurogenesis generated extratelencephalic neuron; RG - radial glia; IP – intermediate progenitor. Scatter points in bar plots indicate individual data points. Error bars indicate SD. Statistical tests used two-sample t-tests (p-values: *p < 0.05, n.s = not significant) Scalebars in (B) 1 mm (C1-C3) 500 µm.

Indeed, retro-orbital co-injection of tTA-driven *AAV-TRE-GFP* and Cre-driven *AAV-FLEX-RFP* in *FezF2-TC;Tbr2-Flp* double transgenic mice resulted in reliable differential labeling of ETs^d^ and ETs^i^ respectively, across the cortex (**Figure 1B**). As negative controls, we verified that *AAV-FLEX-RFP* showed no leaky expression in the cortex (**Figure S1H-I**), and *AAV-TRE-GFP* showed minimal leaky expression that was restricted to somatic but not axonal labelling, even after immunofluorescence amplification (**Figure S1D-G**).

To further validate our strategy, we compared the numbers and relative ratios of AAV-based ET^d^ and ET^i^ labeling with those of a previously validated intersection-subtraction reporter allele (IS)-based labeling ^17^. *AAV-TRE-GFP* and *AAV-FLEX-RFP* co-injection in motor, barrel, and visual cortex in *FezF2-TC;Tbr2Flp* mice yielded very similar labeling of ET^d^ and ET^i^ (**Figure 1C-C3**) in equivalent areas in *FezF2-CreER;Tbr2-Flp;IS2* mice (**Figure S1J-M**). In both approaches, the number of ET^i^ neurons was significantly more than ET^d^ for motor and barrel cortices (**Figure 1K-L**), and there was no significant difference for the visual cortex although there were more ET^i^ neurons (**Figure 1M**). Although viral labeling captured greater number of ET^d^ and ET^i^ neurons in *FezF2-TC;Tbr2Flp* as compared to the IS2 reporter strategy, likely due to higher sensitivity of AAV labelling, there were no significant differences in their proportion, which ranged from 20-30% for ET^d^ (mean: 24.5%) and 70-80%for ET^i^ (mean:75.5%) for all three areas from either approach (**Figure S1N-N2**). Thus, ETs^d^ and ETs^i^ are generated across different cortical areas with an overall ratio of approximately 1:3, indicating differential amplification by iNG. Together, these results validate our novel strategy for differential AAV targeting of ET^d^ and ET^i^ in the same animal.

### Amplification, diversification and innovation of ET projection types by iNG

While ETs^d^ and ETs^i^ are distributed across different regions of the isocortex^17^, whether and how they differ in their projection targets is unknown. We first addressed this question by retrograde labelling from four subcortical targets – the superior colliculus (SC), pontine nuclei (PONS), medulla (Med) and the spinal cord (SpC) (**Figure 2A-D**). Given that the two retro-AAVs (*AAV-TRE-RFP* and *AAV-FLEX-GFP* that label ET^d^ and ET^i^, respectively) are co-injected in the same location of the same animal, this method allows us to compare the distribution and relative proportion of ET^d^ and ET^i^ across the isocortex. Brains were analyzed by both confocal (n = 2, each target area, **Figure 2E-H**) and lightsheet microscopy (n = 2-3 each target area, **Figure 2A-D**, **S2E**, **Video S1-S4**). Indeed, retroAAV injections from the 4 target sites labeled intermixed layer 5 ET^d^ and ET^i^ that were distributed across the isocortex (**Figure 2A-D**, **S2A-D**).

**Figure 2.**
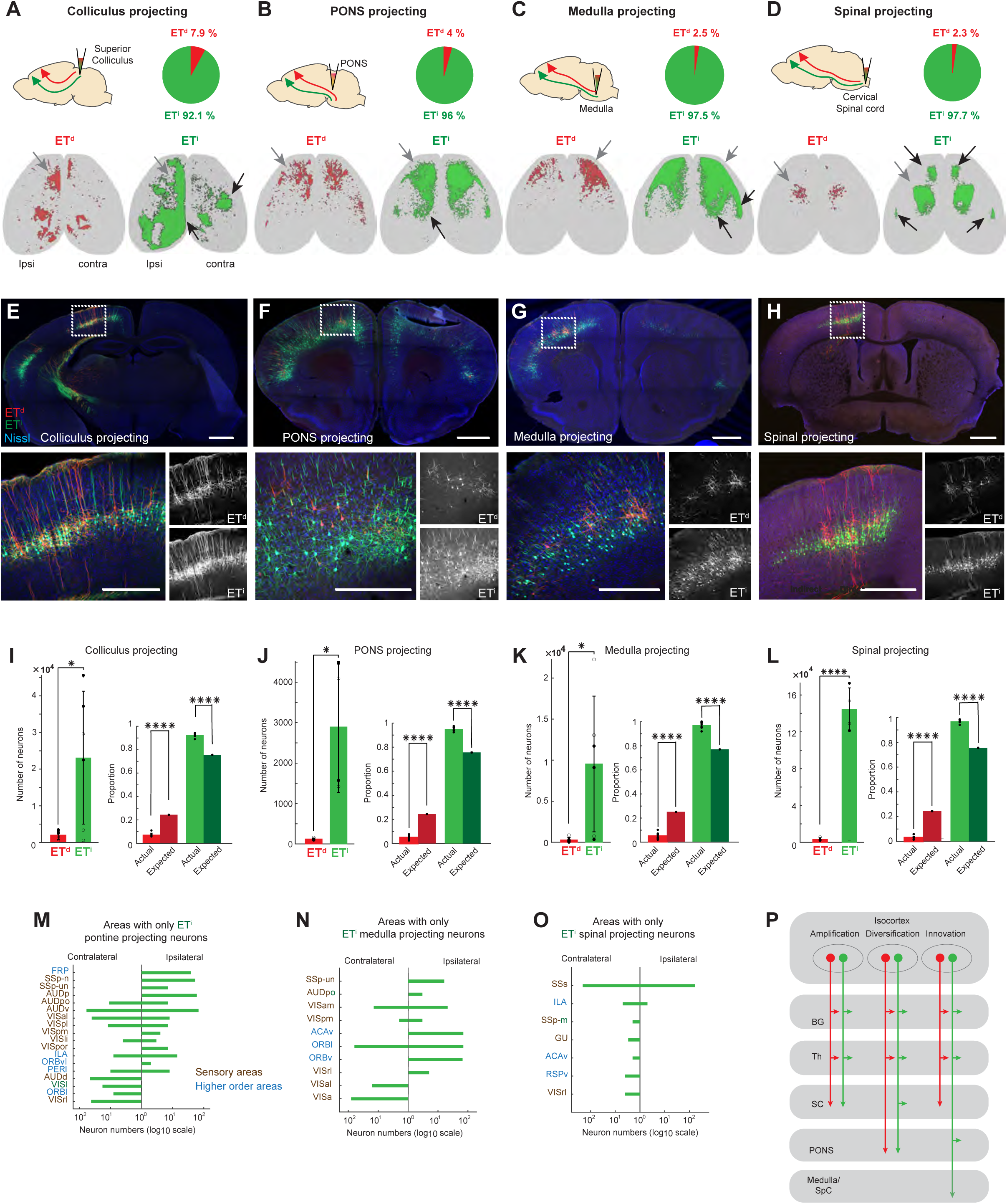
Amplification, diversification and innovation of ET^i^ over ET^d^ subclasses. (A-D) Retrograde labelling of ET^d^ (red) and ET^i^ (green) from superior colliculus (A), pontine nuclei (B), medulla (C), and cervical spinal cord (D). Injection schematics are in top left. Pie charts show the overall proportion of labeled ET^d^ and ET^i^across isocortex for each injection site. Bottom row shows the 3D rendering of the distribution of labelled ET^d^ and ET^i^neurons across isocortex. Green arrows indicate example areas with both ET^d^ and ET^i^; red arrows indicate example areas with significantly more or entirely ET^i^. (E-H) Retrogradely labelled ET^d^ and ET^i^ for (E) colliculus-projecting, (F) pons-projecting, (G) medullary-projecting and (H) spinal-projecting neurons across the isocortex. (I-L) Left panel*s*: Bar plots showing that the number of retrogradely labeled ET^i^ is significantly larger than that of ET^d^across isocortex from each of the four injection sites. Right panels: Paired bar plots comparing the expected proportions of ET^d^ and ET^i^ (based on their approximate overall 1:3 cell number ratio) to the actual labelled proportions for each injection site. There is a significant increase in the proportion of actual ET^i^ over expected ET^i^ for all 4 injection sites. On the other hand, there is a significant decrease of proportion of actual ET^d^ over expected ET^d^. (M-O) Bar plots of retrogradely labelled ET neuron numbers for areas with exclusively ET^i^ and no ET^d^ from pontine (M), medullary (N), and spinal (O) injections, suggesting the innovation of novel ET^i^ over ET^d^ projection types in these areas. Note the abundance of sensory (brown font) and higher order (blue font) areas. There were no areas with exclusively ET^i^ labeling from collicular injection. (P) Schematic representing amplification, diversification and innovation of ET subclasses by iNG. Amplification is simple generation of copies of ET^d^ by ET^i^; Diversification is generation of distinct projection subclasses by ET^i^ while innovation reflects generation of novel projection classes which are exclusively ET^i^. Abbreviations: BG – basal ganglia; Th – thalamus; SC – superior colliculus; PONS – pontine nuclei; SpC – spinal cord. For abbreviations in (M-O) see Table S1. Scatter points in bar plots indicate individual data. Filled markers indicate ipsilateral and open markers indicate contralateral data.Error bars indicate SD. Statistical tests used: two-sample t-tests (p-values: *p < 0.05, ****p < 0.0001, n.s = not significant) Scalebars in (E-H, top panels) 1 mm (E-H, bottom panels) 500 µm.

Considering the overall proportions of ET^i^ and ET^d^ neurons across isocortex, the amplification of ET^i^ cell number over ET^d^ is approximately 3:1 (**Figure S1N-N2**). If ET^i^ and ET^d^ neurons broadly project to the same sets of subcortical targets, we would broadly expect a similar degree of 3:1 *amplification* of ET^i^ over ET^d^ projection at these targets. However, retrograde labeling revealed a substantially disproportionate increase in L5 ET^i^ over ET^d^ proportion from all four injection sites (**Figure 2I-L**): the ET^i^/ ET^d^ ratio ranged from 12.5:1 for collicular-projecting, 25:1 for pontine-projecting, 33:1 for medullary-projecting, to 40:1 for spinal-projecting neurons. As each ET neuron often projects to different combinations of these targets, this result indicates a substantial *diversification* of projection patterns for ET^i^ over ET^d,^, especially those to the more distal targets assessed. Moreover, while both ET^i^ and ET^d^ neurons were mostly co-labeled and intermixed across areas, though at different ratios, by colliculus injection, there were multiple areas (e.g., SSp-n, AUDpo, ORBv, ILA) where only ET^i^ but not ET^d^ neurons were labeled by injection to the PONS, Med, and SpC (**Figure 2M-O**). This result indicates the presence of novel distally-projecting ET types in these areas generated only through iNG, which we consider as *innovation* by iNG (**Figure 2P**). Notably, these novel area-specific ET^i^ projection types were largely found in sensory and higher order areas rather than motor areas (**Figure 2M-O**). This result suggests that in addition to the diversification of ET projection types from the frontal and motor areas, iNG generates novel projection types in some non-motor areas that grant them direct access to the motor centers and other areas in the PONS, Med and SpC.

Although the 4 injection sites represent only a small subset of ET targets, the results together demonstrate that, compared with dNG, iNG not only amplifies but also diversifies projection classes, and additionally generates novel ET projection types. To reveal the precise and comprehensive projection patterns of ET^d^ and ET^i^ in specific areas, we then carried out a set of anterograde labeling across eight isocortical areas encompassing motor, sensory, and higher order areas.

### ET^i^ in motor areas expands projections to action diversification and execution circuits

For the motor areas, we chose the rostral forelimb orofacial area (RFO ^33^) and the caudal forelimb area (CFA^11,34–36)^ in the primary motor cortex (MOp) and an area in the secondary motor cortex (MOs) recently implicated in a forelimb-reach-to-grasp behavior (RGD) ^37^(**Figure 3A-C**, **S3A, S4A, S5A**). In all three areas, *AAV-TRE-GFP* and *AAV-FLEX-RFP* labelled ET^d^ and ET^i^ in deep layers, with approximately 1:3 ratio (**Figure S3A1-A2, S4A1-A2, S5A1-A2**). Our goal here is to quantitatively compare the relative strength of ET^i^ and ET^d^ projections across major subcortical targets. While confocal microscopy gave high resolution view of subsets of targets (n = 2 for each area, **Figures S3A-G, S4A-G, S5A-G**), light-sheet microscopy enabled us to capture the entire projections to all targets in the whole brain (n = 2 for each area, **Figure 2A-C, S3I, S4I, S5I, Video S5-S7**). We generated 3D renderings of projections and fraction occupation heatmaps for all downstream targets (listed in **Table S1**) for the 3 injection areas (**Figure 3A-C, 3D-F**).

**Figure 3.**
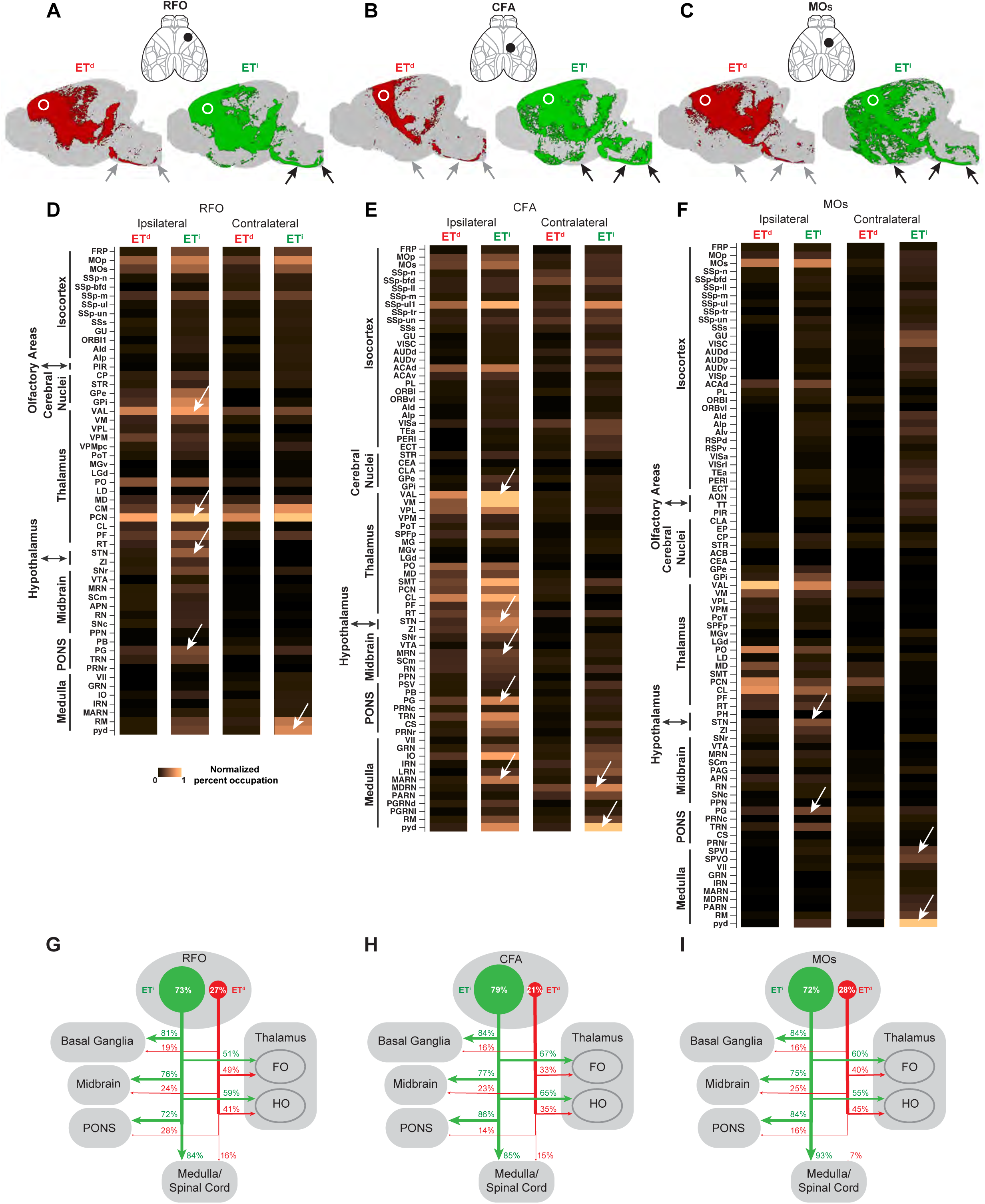
Amplification and diversification of ET^i^ over ET^d^ projections from motor areas. (A-C). Schematic of anterograde AAV injection site and 3D rendering in the sagittal plane for ET^d^ and ET^i^ axonal projections from (RFO, A), (CFA, B) and (MOs, C). White circles indicate injection sites. Black arrows indicate examples of greatly expanded ET^i^ projections to more distal targets. (D-F). Heatmaps showing normalized percent occupation of target structures across the brain from ET^d^ and ET^i^ for ipsilateral and contralateral sides of the injection sites for RFO (D) CFA (E) MOs (F). White arrows indicate example areas with significant increase in percent occupation by ET^i^ over ET^d^. Note the substantial increase of ET^i^ projections to several brainstem areas and the pyramidal tract. (G-I). Schematics showing the amplification and diversification of ET^i^ over ET^d^ neurons and projections from the three motor areas scaled to the proportion of ET^d^ and ET^i^ labelled at the injection sites. Note the significantly expanded ET^i^ projections to the BG (action selection), and medulla and SpC (action diversification and execution) areas while having similar projections as ET^d^ to thalamus. Abbreviations: RFO – rostral forelimb orofacial area in the motor cortex; CFA – caudal forelimb area; MOs – secondary motor area; BG – basal ganglia; SpC – spinal cord. For abbreviations in (D-F) see Table S1.

To quantify the relative projection strength of ET^i^ and ET^d^ using light-sheet microscopy datasets, we measured the percent occupancy in selected target structures using ET^d^ or ET^i^ axon fluorescence signals. This measurement is not in terms of fluorescence intensity, which is not directly comparable between tdTomato (ET^i^) and GFP (ET^d^), and could be an underestimate of ET^i^ projection strength. An equivalent ratio of ET^i^ over ET^d^ projection strength to that of their cell body ratio at injection site (e.g., 3:1) would suggest a simple amplification by ET^i^. On the other hand, a disproportional increase of ET^i^ proportions at different target regions would suggest a diversification or innovation of ET^i^ projections. Thus, in addition to assessing the relative increase of ET^i^ over ET^d^ projections at each target, we compared them across different target regions of interest and ran post-hoc comparisons between these regions with the thalamus to reveal specific target regions with a disproportional increase of ET^i^ projections. The thalamus was chosen for comparison as a forebrain region which receives substantial ET^d^ as well as ET^i^ projections.

While both ET^d^ and ET^i^ targeted the thalamus (**Figure S3B, S4B, S5B**), ET^i^ had significantly more projections from all three areas (**Figure S3B1-B3, S4B1-B3, S5B1-B3**). Notably, *FezF2* expressing ET neurons include both PT (pyramidal-tract like, mostly in L5b but also L6) and some CT (cortico-thalamic in L6) populations; thus, the labeled thalamic projections derive from both populations (See Discussion, **Figure S3B, S4B, S5B**).

As a broad category, projections to the brainstem areas (i.e. PT type projections that include MB, PONS, Med, SpC) from all three motor areas were significantly amplified in ET^i^ over ET^d^ (**Figure S3B1-B3, S4B1-B3, S5B1-B3**). Within the thalamus, the three motor areas showed differences in ET^d^ vs ET^i^ projection to first order (FO) and higher order (HO) nuclei. RFO and MOs had similar ET^d^ and ET^i^ projections to FO but more ET^i^ projections to HO (**Figure S3C1-C3, S5C1-C3**). CFA had stronger ET^i^ projections to both FO and HO (**Figure S4C1-C3**).

At the level of basal ganglia (BG), ET^i^ had a major enhancement of projections over ET^d^ to all BG nuclei from all three areas (**Figure S3D1-D3, S4D1-D3, S5D1-D3, Figure 3G-I**) indicating vastly expanded control by ET^i^ over action selection circuits ^38,39^.

In the MB, ET^i^ had significantly more projections than ET^d^ from all three areas (**Figure S3E1-E3, S4E1-E3, S5E1-E3**); this was also true for projections to the PONS and Med (**Figure S3F1-F3, S3G1-G3, S4F1-F3, S4G1-G3, S5F1-F3, S5G1-G3)**. When the relative increase of ET^i^ over ET^d^ was compared across target regions, there were significant differences (p value for RFO:p < 0.05; CFA:p < 0.0001; MOs:p < 0.01, Kruskal–Wallis test). Post-hoc comparisons revealed that for all three areas, increase of ET^i^ over ET^d^ to the Med and BG and additionally to PONS were more pronounced compared to that in thalamus (**Figure S3H, S4H, S5H**). This result suggests that Med-projecting ET from RFO and PONS-and Med- projecting ET from the CFA and MOs are largely generated by iNG.

Thus, across motor areas, while both ET^d^ and ET^i^ target forebrain and midbrain areas, the bulk of corticofugal projections to more distal targets are predominantly generated by iNG-ET^i^ to innervate action selection, diversification and execution centers in the BG, hindbrain and SpC, (**Figure 3G-I**). This expansion by iNG-ET^i^ thus creates the anatomical substrate for monosynaptic control of motor centers in the BS and SpC by motor areas of the isocortex.

### ET^i^ in sensory areas expands subcortical projections to ascending sensory pathways

We targeted three sensory areas for anterograde tracing – the visual cortex (VIS), auditory cortex (AUD) and barrel cortex (BAR) (n=2 [confocal, **Figure S6A-E, S7A-G, S8A-G**], n = 2 [lightsheet, **Figure S6G, S7I, S8I, Video S8-S10**] for each area, **Figure 4A-C *top*, S6A, S7A, S8A**). The VIS injection labelled neurons in layers 5 and 6 with more ET^i^ (**Figure S6A1-A2, S7A1-A2, S8A1-A2**). In AUD and BAR areas, there were more ET^i^ labelled in layer 5; while in layer 6a/6b, more ET^d^ or near equal numbers of ET^d^ and ET^i^ were labelled (**Figure S7A1-A2, S8A1-A2**). The larger number of ET^d^ in layer 6a/6b are likely CT neurons, which are mostly derived from dNG^17^. We generated 3D renderings of projections and heatmaps for all three injection areas (**Figure 4A-C, 4D-F**).

**Figure 4.**
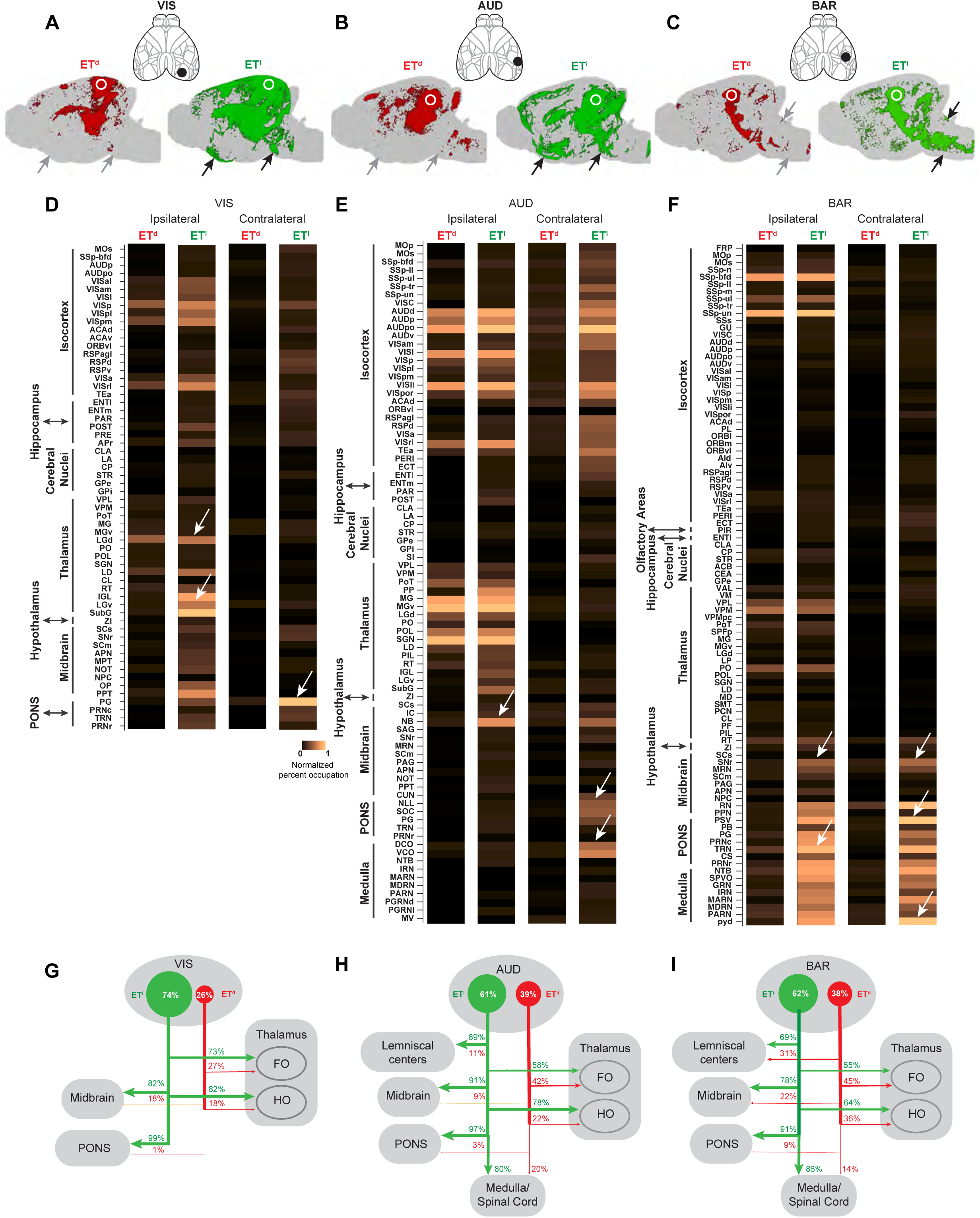
Amplification, diversification and innovation of ET^i^ over ET^d^ projections from sensory areas. (A-C). Schematic of anterograde AAV injection site and 3D rendering in the sagittal plane for ET^d^ and ET^i^ axonal projections from (VIS, A), (AUD, B) and (BAR, C). White circles indicate injection sites. Black arrows indicate examples of greatly expanded ET^i^ projections to more distal targets. (D-F). Heatmaps showing normalized percent occupation of target structures across the brain from ET^d^ and ET^i^ for both ipsilateral and contralateral sides of the injection sites for VIS (D) AUD (E), and BAR (D). White arrows indicate example areas with significant increase in percent occupation by ET^i^ over ET^d^. Note the increased or exclusive projections to PONS from ET^i^ in all three areas while similar increased projections to medulla from ET^i^ from BAR. (G-I). Schematics showing the amplification and diversification of ET^i^ over ET^d^ neurons and projections from the three sensory areas scaled to the proportion of ET^d^ and ET^i^ labelled at the injection sites. Note that the corticopontine projections are almost exclusively ET^i^ projections and the massive ET^i^ elaboration to lemniscal centers. There is also an expansion of ET^i^ projections to medulla and SpC, similar to motor areas from BAR. Abbreviations: VIS – visual cortex; AUD – auditory cortex; BAR – barrel cortex; PONS – pontine nuclei; SpC – spinal cord. For abbreviations in (D-F) see Table S1.

From VIS and AUD, there was significantly more ET^i^ than ET^d^ projections to both the thalamus and brainstem areas (**Figure S6B1-B3, S7B1-B3, S8B1-B3**). Within the thalamus, there was no significant difference between ET^d^ and ET^i^ projections to FO while ET^i^ had significantly more projections to HO (**Figure S6C1-C3, S7C1-C3, S8C1-C3**). The dense projections of ET^d^ to FO may include a significant CT component while those of ET^i^ to both FO and HO could largely derive from PT.

For AUD and BAR areas, we compared ET projections to major sensory processing stations along their respective ascending lemniscal pathways. ET^i^ but not ET^d^ heavily targeted lemniscal centers, suggesting that iNG confers sensory cortical access to lemniscal sensory processing centers (**Figure S7D1-D3, S8D1-D3**).

When examining VIS corticofugal projections, we noticed that in some prethalamic nuclei such as the ventral lateral geniculate (LGv), the ET projections were almost entirely ET^i^ (**Figure S6C**). Notably, LGv is expanded in nocturnal mammals such as rodents and is similar in size to the LGd ^40^.

Regarding ET projections to the MB, projections from all three areas were dominated by ET^i^ (**Figure S6D1-D3, S7E1-E3, S8E1-E3**). There were generally more ET^i^ than ET^d^ projections to PONS and Med (**Figure S6E1-E3, S7F1-F3, S8G1-G3, S8F1-F3, S8G1-G3**). Notably, the relative increase of ET^i^ over ET^d^ for VIS was not different between target regions indicating a similar increase of ET^i^ projections across subcortical structures (**Figure S6F**). In contrast, the increase of ET^i^ over ET^d^ from AUD for the lemniscal centers and PONS was larger than that in the thalamus (**Figure S7H**); this was true across the BS for BAR (**Figure S8H**).

Thus, in sensory areas, iNG-ET^i^ elaborates projections to HO thalamus, prethalamus and sensory lemniscal centers, while also incorporating motor like features to BAR with elaborated projections to hindbrain. In contrast, ET^d^ heavily targets FO thalamus, with projections also to the midbrain and weak projections to lemniscal centers, PONS and Med (**Figure 4G-I**).

### ET^i^ in higher order areas diversifies projections to pallial, hypothalamic and neuromodulatory structures

We further compared ET^i^ and ET^d^ projections in two higher order areas of the isocortex, the prefrontal (PFC) and posterior parietal areas (PAR) (n=2 for confocal, [**Figure S9A-J, S10A-G**], n = 2 for Lightsheet, **Figure 5A-B, S9L,S10I,Video S11-S12** ] for each area. The injection at each area labelled neurons with more ET^i^ than ET^d^ (**Figure S9A-A2, S10A-A2**).

**Figure 5.**
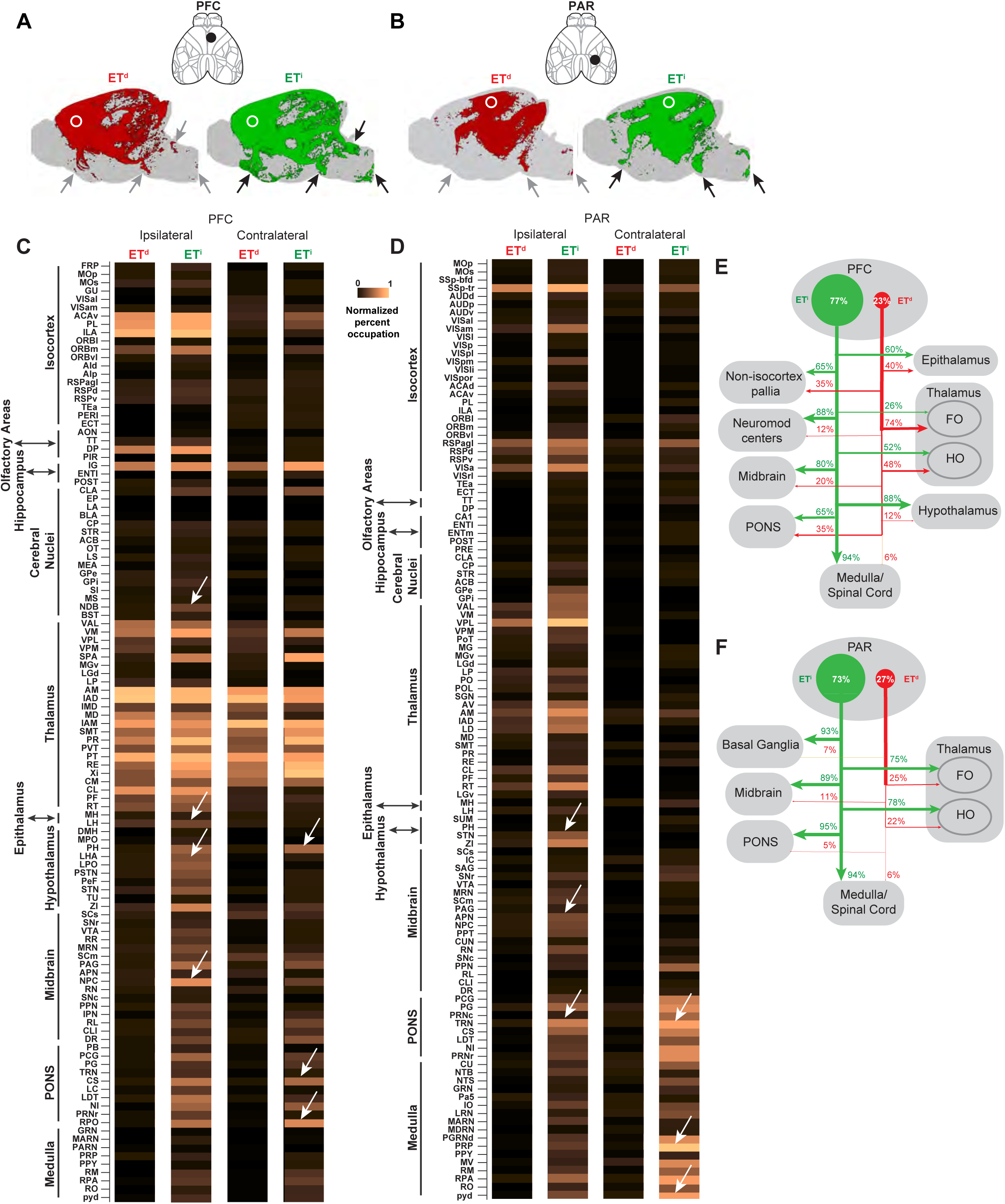
Amplification, diversification and innovation of ET^i^ over ET^d^ projections from higher order areas. (A-B). Schematic of AAV injection site and 3D renderings in the sagittal plane for ET^d^ and ET^i^ axonal projections from (PFC, A) and (PAR, B). White circles indicate injection sites. Black arrows indicate examples of greatly expanded ET^i^ projections to more distal targets. (C-D). Heatmaps showing normalized percent occupation of target structures across the brain from ET^d^ and ET^i^ for both ipsilateral and contralateral sides of the injection sites for PFC (C) and PAR (D). White arrows indicate example areas with significant increase in percent occupation by ET^i^ over ET^d^. Note the significantly increased ET^i^ projections to hypothalamus, other pallial areas and the brainstem, while similar projections as ET^d^ in the thalamus from PFC. For PAR note the increased ET^i^ projections to BG and brainstem areas. (E-F). Schematics showing the amplification and diversification of ET^i^ over ET^d^ neurons and projections from PFC and PAR scaled to the proportion of ET^d^ and ET^i^ labelled at the injection sites. Note the expansion of ET^i^ projections to neuromodulatory, hypothalamic and other pallial areas while having similar proportions as ET^d^ to the epithalamus from PFC. Note motor-like expansion of ET^i^ projections to BG and medulla/SpC from PAR. Abbreviations: PFC – prefrontal cortex; PAR – parietal cortex; BG – basal ganglia; SpC – spinal cord. For abbreviations in (C-D) see Table S1.

From the PFC, both ET^i^ and ET^d^ targeted the thalamus heavily while there were stronger ET^i^ projections over ET^d^ to the brainstem regions (**Figure S9B1-B3**). In contrast, from PAR,there were more ET^i^ projections over ET^d^ to both thalamus and BS, similar to what was observed in motor areas (**Figure S10B1-B3**). This result suggests higher numbers of PT neurons in PAR sending collaterals to thalamus.

Comparing projections to FO and HO thalamus, from PFC ET^d^ had stronger projection to FO than ET^i^, while ET^i^ had stronger projection to HO. From PAR, ET^i^ had stronger projections to both FO and HO (**Figure S9C1-C3, S10C1-C3**). The PAR projections to FO and HO were similar to proportions of ET^i^ and ET^d^ projections from motor areas (**Figure S10C1-C3**). Given this similarity and since the PAR is known to have roles in motor behavior^41,42^, we further examined BG and found again that the projections were predominantly ET^i^ **(Figure S10D-D3)**.

The PFC is known to target multiple non-isocortical pallial areas ^43^. We found that the majority of these projections were ET^i^ (**Figure S9D1-D3**), suggesting a massive elaboration of inter-pallial interaction by iNG- ET^i^. Some other forebrain areas like the epithalamus received similar PFC ET^d^ and ET^i^ projections (**Figure S9E1-E3**). Notably, ET^i^ projections dominated the hypothalamus (**Figure S9F1-F3**). This highlights selective expansion of projections by distinct neurogenic pathways in anatomically/developmentally close and related structures.

Another important observation was the heavy ET^i^ over ET^d^ projections to different neuromodulatory centers across the brain from PFC (**Figure S9G1-G3**), indicating the expansion of higher order cortical control of neuromodulation. PFC and PAR projections to the MB and PONS were largely ET^i^ (**Figure S9H-H3, S9I-I3, S10J-J3, S10E-E3, S10F-F3, S10G-G3**).

Taken together and considering the relative increase of ET^i^ over ET^d^ among different target regions and the thalamus, ET^i^ in higher order areas selectively expands projections to neuromodulatory centers, hypothalamus and also dominates interactions with other pallial areas while incorporating motor area like features with significant expansions to hindbrain and SpC (**Figure S9K, S10H, Figure 5E-F**).

### Selective axon pruning in ET^i^ but not ET^d^ during early postnatal development shapes the areal distribution of ET^i^

The area specific ET projection patterns to multiple subcortical regions are established by a multi-step developmental process that involves progressive (axon guidance, synapse formation) as well as regressive (axon pruning, synapse elimination) events ^44,45^. In particular, axon pruning has been demonstrated for corticofugal neurons from multiple cortical areas during early postnatal period ^46^, with a prominent role in shaping the arealization of descending pathways ^47^. We therefore examined whether axon pruning may differentially sculpt the projection patterns and arealization of ET^d^ and ET^i^. As substantial pruning of mouse corticospinal axons occurs during the first postnatal week, we injected retroAAVs in the cervical spinal cord (cSpC) of P4 mice and examined cortex-wide ET^d^ and ET^i^ labelling pattern at P40 (P4-P40), and compared it with those labeled from adult (P28) cSpC injection (P28-P40) (n = 2, confocal, **Figure 6C-D**; n = 2, lightsheet, **Figure 6A-B, S11D, Video S13**).

**Figure 6.**
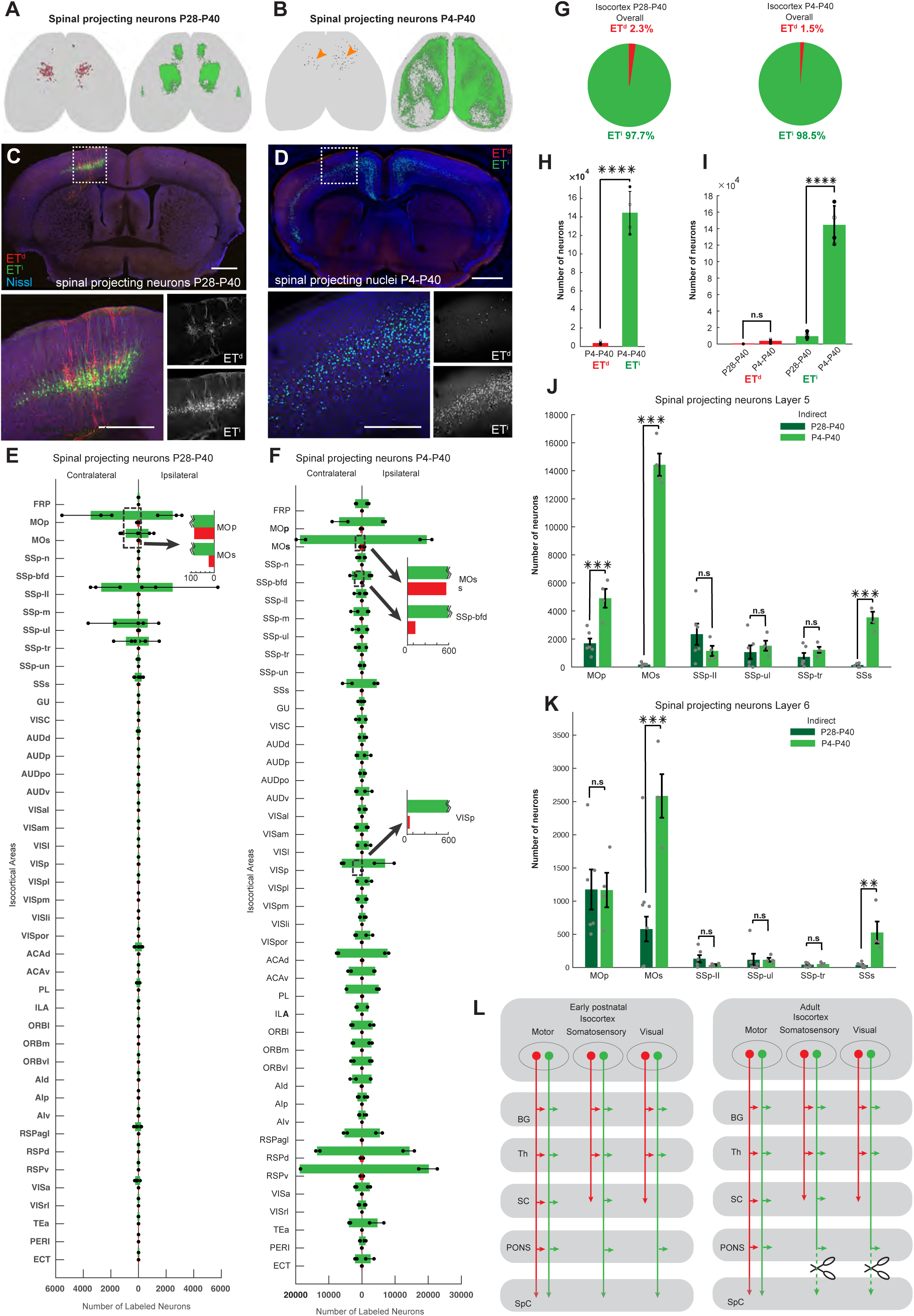
Selective axon pruning in ET^i^ but not ET^d^ neuron during early postnatal development shapes the areal distribution of ET^i^. (A-B) 3D renderings of retrogradely labelled ET^d^ (red) and ET^i^ (green) spinal projecting neurons/nuclei across isocortex from retrograde AAV injections into the cervical spinal cord of P28-P40 (A) and P4-P40 (B) mice. (C-D) Retrogradely labelled ET^d^ and ET^i^ spinal projecting neurons (C) or nuclei (D) from cSpC injections in P28-P40 (C) and P4-P40 (D) mice. Scale bars: 1 mm in top and 500um in bottom panels. (E-F) Bar plots showing number of retrogradely labelled ET^d^ (red) and ET^i^ (green) spinal projecting neurons across different isocortical areas. Error bars represent standard deviations. Zoomed in views magnify selected areas (in dotted black rectangle) showing low ET^d^ numbers compared to ET^i^. The zoomed ET^i^ bar plots are truncated to show ET^d^ cell numbers. (G) Pie charts showing the proportion of ET^d^ and ET^i^ spinal projecting neurons across isocortex from injections into the cervical spinal cord of P28-P40 (left) P4-P40 (right) mice. (H) Bar plots showing that the number of retrogradely labelled ET^i^ spinal projecting neurons is significantly larger than the number of ET^d^ spinal projecting neurons across isocortex from injections into cSpC of P4-P40 mice. (I) Bar plots showing that number of ET^i^ spinal projecting neurons across isocortex from cSpC injections in P4-P40 mice is significantly more than that of cSpC injections in P28-P40 mice, while there is no significant change for ET^d^spinal projecting neurons between the two. (J) Paired bar plots comparing number of retrogradely labelled layer 5 spinal projecting neurons/nuclei in MOp, MOs, SSp-ll, SSp-ul, SSp-tr and SSs between P28-P40 and P4-P40 mice. The P4-P40 mice had significantly more nuclei in MOp, MOs and SSs while there were no significant changes in other areas. (K) Paired bar plots comparing number of retrogradely labelled layer 6 spinal projecting neurons/nuclei in MOp, MOs, SSp-ll, SSp-ul, SSp-tr and SSs between P28-P40 and P4-P40 mice. The P4-P40 mice had significantly more nuclei in MOs and SSs while there were no significant changes in other areas. **(O)** A schematic showing selective axon pruning in corticospinal ET^i^ but not ET^d^ neurons during early postnatal development more evident in mid and posterior areas of the isocortex, which shapes the projection patterns and areal distribution of ET^i^ in mature cortex. Abbreviations: P28-P40 – mice with injections on P28 and harvested at P40; P4-P40 – mice with injections at P4 and harvested at P40; cSpC – cervical spinal cord. For abbreviations in (E-F, J-K) see Table S1. Error bars indicate SD. Scatter points indicate individual data.Filled markers indicate ipsilateral and open markers indicate contralateral data Statistical tests used: (H, J, K) two-sample t-test; (I) two-sample t-test and Mann-Whitney U test (p-values: *p < 0.05, **p < 0.01, ***p < 0.001,****p < 0.0001, n.s = not significant)

In contrast to the P28-P40 ET labeling pattern that is highly restricted to a few areas (MOp, MOs, SSp, **Figure 6E**), P4 cSpC injection labeled a significantly larger number of ET neurons across the anterior-posterior extent of cortex (**Figure 6F**), consistent with previous findings^47,46^. Strikingly, this vast expansion of P4-P40 ET labeling consisted of almost exclusively ET^i^ (**Figure 6G-I**). In contrast, P4-P40 labeled a small number of ET^d^ neurons that remained restricted to the same few areas as those in P28-P40 labeling (**Figure 6E-I**). This result suggests that across the early postnatal cortex, ET^i^, but not ET^d^, initially undergo exuberant axon growth to the spinal cord; and the massive cortex-wide pruning of ET neuron axons that sculpts their subsequent areal specific patterns is restricted to ET^i^. In contrast, most ET^d^ neurons across the developing cortex do not project to the spinal cord, and the mature pattern of ET^d^-spinal projection in specific areas (e.g. MOp, MOs and SSp) is likely achieved by area-specific properties of axon growth and guidance.

Notably, ET neurons reside in both L5 and L6, and many L6 ET neurons project to more proximal subcortical targets than the spinal cord. In this context, we found that P4 cSpC injection labeled many L6 ET^i^ but not ET^d^ in MOs and SSs (**Figure 6J-K**). This result suggests that the corticofugal projection pattern of L6 ET^i^, but not ET^d^, is also shaped by the pruning of their axon arbors that initially reached the spinal cord.

Surprisingly, we found that P4 cSpC injection further labeled iNG- but not dNG-derived neurons in many pallial areas outside of the isocortex (**Figure S11A-C, S11E-G**). The largest number were found in the hippocampal formation (**Figure S11C, S11E**) but also in other regions including piriform cortex (**Figure S11A**) and amygdala (**Figure S11B**). As controls, cSpC injection at P1, when ET axons have not reached the spinal cord, resulted in very sparse labelling of ET^d^ and ET^i^ in cortical structures at P40 (n = 2, **Figure S11G**). Further, sSpC injection of a generic retro-AAV into wild type mice at P6 labeled neurons in both the isocortex and other pallial areas, confirming our results using *FezF2TCTbr2Flp* mice and Cre and tTA dependent retro-AAvs (n = 2, **Figure S11I**). All together, these results suggest that an initial exuberant axon growth, many reaching the SpC, followed by extensive axon pruning, is a general feature of ET^i^ but not ET^d^ across the pallial/cortical structures to establish their areal specific subcortical projection patterns.

### Functional distinctions of ET^d^ and ET^i^ in the motor cortex

To uncover the functional significance of the projection differences between ET^d^ and ET^i^, we carried out differential optogenetic stimulation of ET^d^ and ET^i^ in the same animal in head-fixed as well as free-moving conditions using *the FezF2-TC-Tbr2 mice* (**Figure 7A,7C,7J-J3**). We injected *AAV-TRE-ChRmine-oScarlet* on the left RFO and *AAV-FLEX-ChRmine-oScarlet* on the right RFO (**Figure 7C**). The injections robustly captured ET^d^ and ET^i^ (**Figure 7C**) which could be optogenetically activated sequentially in the same mouse. As a control, we also performed similar activation of all ET neurons (ET^all^) using *FezF2-CreER;Ai32* mice^33,48^ (**Figure 7B)**.

**Figure 7.**
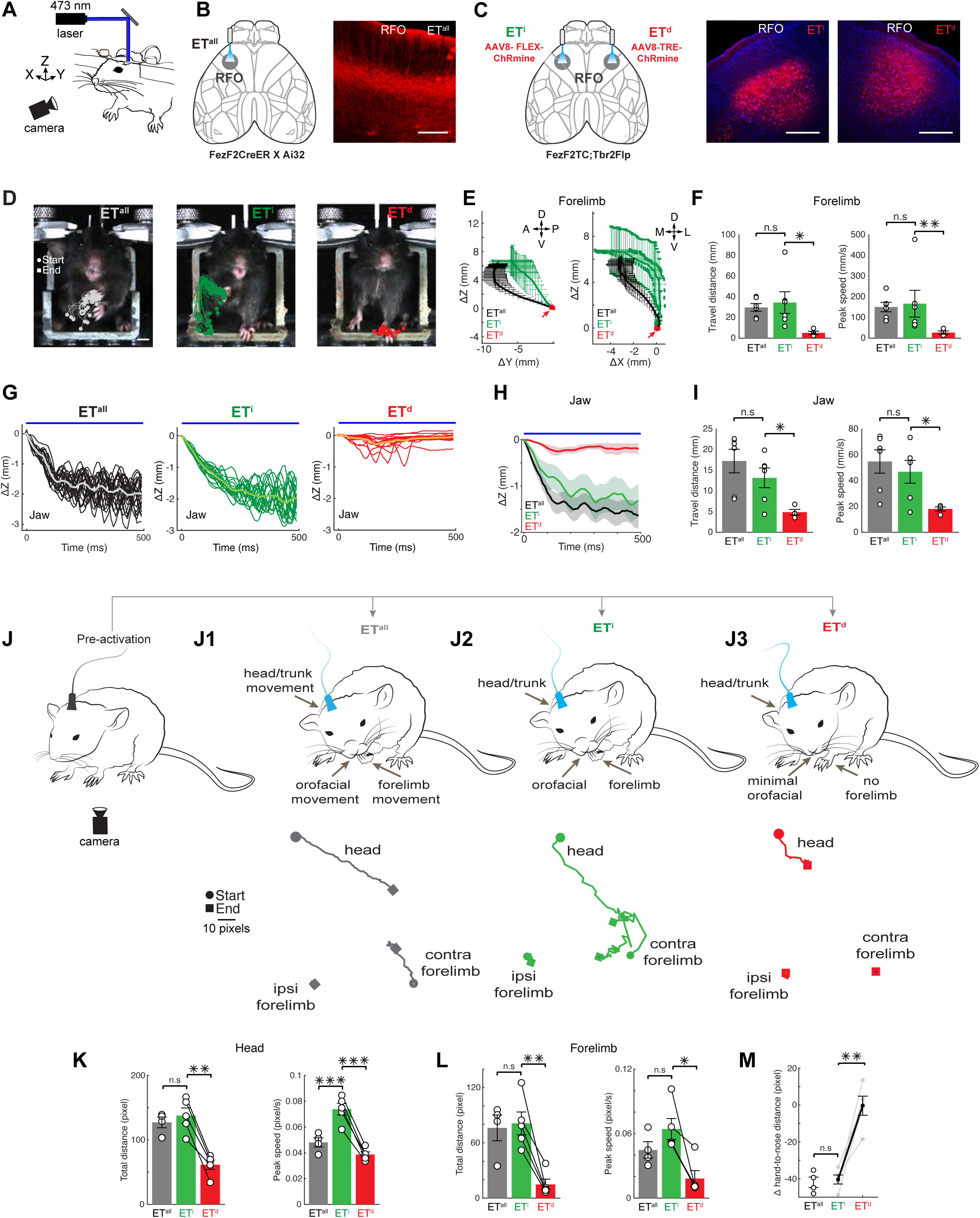
Optogenetic activation of ET^d^ and ET^i^ in the motor cortex reveal distinct roles in controlling movement. (A-C) Schematic showing experimental setup for head-fixed optogenetic activation (B) Schematic showing activation of ET^all^ in the RFO in *Fezf2CreERAi32* mice; ET^all^ neurons expressing channelrhodopsin (right). (B) Schematic showing activation ET^i^ and ET^d^ in the same *Fezf2Tbr2flp* miceRFO (left). ET^d^ is targeted with AAV8-TRE-ChRmine-oScarlet on the right RFO while ET^i^ is targeted with AAV8-FLEX-ChRmine-oScarlet on the left RFO.Labelling of ET^i^ and ET^d^ (right) showing expression of ChRmine (magenta) in the left and right RFO respectively. (D) Contralateral forelimb trajectories during activation of ET^all^ (20 trials, left), ET^i^ (20 trials, middle) and ET^d^ (17 trials, right) in RFO with circles and squares indicating start and end points respectively. Thick line indicates the average trajectory. Note that forelimb movement was absent during ET^d^ activation. (E) 2D projections of forelimb movement trajectories during optogenetic activation of ET^all^ (n = 6) and ET^i^ (n = 6) and ET^d^ (n = 4) . Thick line indicates the average position. Movement directions are indicated as: A, anterior; P, posterior; D, dorsal; V, ventral; M, medial; L, lateral. (F) Bar plots showing no significant differences in distance travelled (left) and peak speed (right) upon optogenetic activation for the forelimb between ET^all^ (n = 6) and ET^i^ (n = 6) and significant differences between ET^d^ (n = 4) and ET^i^ (n = 6). (G-H) Jaw movement trajectories during optogenetic activation (top blue bar) of ET^all^ (20 trials, left), ET^i^ (20 trials, middle) and ET^d^ (17 trials, right) and (H) across different mice for ET^all^ (n = 6), ET^i^ ( n = 6) and ET^d^ (n = 4). (I) Bar plots showing no significant differences in distance travelled (left) and peak speed (right) upon optogenetic activation for the jaw between ET^all^ (n = 6) and ET^i^ (n = 6) and significant differences between ET^d^ ( n = 4) and ET^i^ (n = 6). (J-J3) Illustrative schematics with corresponding movement trajectory plots of nose and contra/ipsi forelimb showing behavioural effects of optogenetic activation in free moving mice (single trial) in the RFO (preactivation, J) for ET^all^ (J1), ET^i^ (J2) and ET^d^ (J3). Note the significantly amplified nose movement and unique contralateral forelimb movement upon ET^i^ activation. (K-L) Bar plots showing significant differences in distance travelled (left) and peak speed (right) upon optogenetic stimulation in free-moving mice for the nose (K) and forelimb (L) for ET^all^ (n = 4), ET^i^ (n = 5) and ET^d^ (n = 5). (M) Plot showing change in hand to nose distance upon optogenetic activation of ET^all^ (n = 4), ET^i^ (n = 5) and ET^d^ (n = 5) in free-moving mice, with significantly higher changes in ET^i^ while that of ET^d^ is almost zero. Abbreviations: RFO - Rostral forelimb orofacial area; Error-bars indicate SEM. Statistical tests used: Two-sided paired t-test or Two-sided Wilcoxon signed-rank test. p-values: *p < 0.05, **p < 0.01, ***p < 0.001. Movement trajectories were normalized to the start position in (D) and (E).Scalebars in (B) 500 µm, (C) 5 mm.

In the rostral forelimb-orofacial area (RFO), activation of ET^all^ in the head-fixed condition induces contralateral forelimb adduction to the body midline with hand supination and digit closing and jaw opening (**Figure 7D-I; Video S14;** also see^34^). Strikingly, sequential optogenetic activation of ET^d^ and ET^i^ in the left and right RFO in head-fixed condition induced highly distinct movements. Whereas ET^d^ activation induced only minor or no oral movement and no forelimb movements (**Figure 7D-I**; **Video S14; ET^d^:n = 4)**, ET^i^ activation induced prominent and coordinated forelimb and oral movements (**Figure 7D-I**; **Video S14; ET^i^: n = 6**). These movements elicited upon ET^i^ activation resembled that of ET^all^ activation (**Figure 7D-I; Video S14; ET^all^:n = 6**)

In the free-moving condition, ET^all^ activation in the RFO induced movements which involve a shoulder adduction that raises the contralateral hand toward the body midline with associated hand supination and digit flexion. In addition, a concurrent ipsiversive head turning and lowering, and bringing the snout toward the raised hand is induced (**Figure 7J-J3; Video S14;ET^all^:n = 4;** also see^34^). These movements resemble ethological feeding behaviour^33^. Interestingly, ET^d^ stimulation in RFO induced head lowering and associated body/trunk movement but with no forelimb movement. In sharp contrast, RFO ET^i^ stimulation induced the fully coordinated head-to-hand as well as hand-to-mouth movements that are similar to those induced by ET^all^ stimulation and resemble ethological feeding **(Figure 7J-M; Video S14;ET^d^:n = 5;ET^i^: n = 5)**.

Together, these results suggest that ET^d^ mainly regulate midline axis movements involving proximal body parts such as head, trunk consistent with their projection targets, while ET^i^ additionally and directly control distal body parts such as the forelimb and hand, thereby coordinating complex and dexterous movements. These data thus show that ET^i^ amplify and diversify ET^d^ motor cortex functions, consistent with their projection differences to brainstem and spinal to action diversification and execution circuits that mediate oral-manual movements.

## Discussion

The mammalian cerebral cortex contains a vast neural network comprising dozens of functional areas, which together integrate sensory information with cognitive and internal state signals to orchestrate behavior. The computational outcome of the cortex is broadcast to multiple subcortical regions by diverse subpopulations of ET neurons across different cortical areas. Decades of studies demonstrate that ET projection patterns vary substantially within as well as across cortical areas and across mammalian species ^5,8–10,49,50^. It has remained an enigma whether this daunting and seemingly unresolvable heterogeneity is stochastic, reflects a continuum, or can be meaningfully classified into ground truth subpopulations relevant to the developmental process and systems level organization and function. Here we dissect the diversity and organization of ET neurons based on a fundamental distinction in their developmental origin – dNG and iNG, which differ not only in progenitor types (RG vs IP) and cell division patterns (asymmetric vs symmetric) but also in evolutionary history. A key experimental entry point is our design of a genetic strategy that converts developmentally transient lineage signals in neurogenesis to permanent “genetic handles” that allow differential viral targeting of postmitotic ET^d^ and ET^i^ in the same individual mice. We found that ET^d^ projections are largely restricted to forebrain and midbrain structures, while ET^i^ differentially amplifies and diversifies these projections in an area-specific way across brain systems, and overwhelmingly dominates the innervation of hindbrain and spinal cord (**Figure 8**). Notably, corticofugal subpopulations in multiple areas are derived from only ET^i^ but not ET^d^, indicating the generation of novel projection types by iNG. Furthermore, areal-specific ET^i^, but not ET^d^, spinal projection pattern is sculpted from the exuberant overgrowth and subsequent massive pruning of an initial cortex-wide population during early postnatal period. Consistent with their differences in projection patterns, optogenetic activation of ET^d^ and ET^i^ in the motor cortex reveal corresponding functional distinctions: while ET^d^ induced movements of head and trunk, ET^i^ additionally elicited coordinated orofacial and forelimb movements that resemble feeding. Together, these results uncover that two foundational neurogenic pathways with distinct evolutionary history differentially shape the area-specific organization and functional diversification of cortical output channels (**Figure 8**).

**Figure 8.**
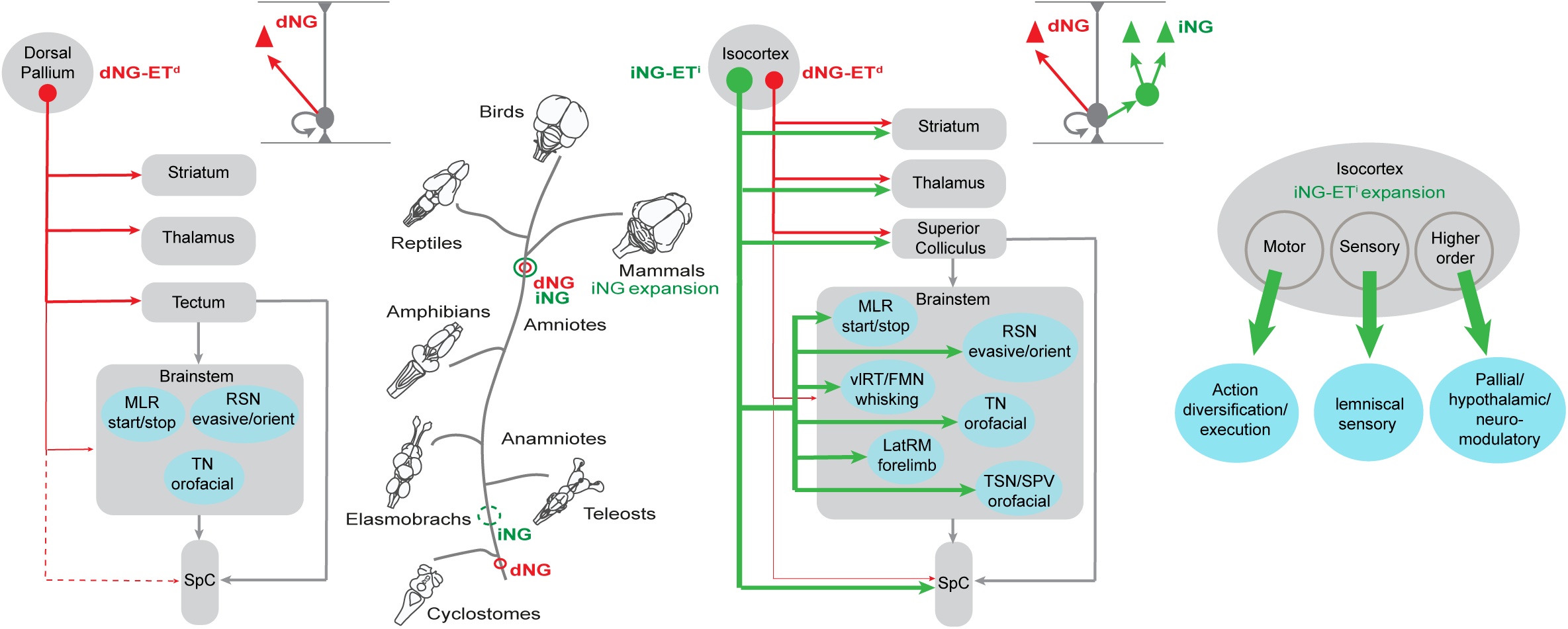
Schematic summary of the evolutionary origin and development impact of dNG and iNG that shape the amplification, diversification and innovation of corticofugal projection neurons. While dNG evolved before the dawn of vertebrates, iNG emerged from amniotes and greatly expanded in mammals. In early vertebrates (left), the pallium does have a basic dNG-ET^d^ (red) efferent projection pattern from the pallium to downstream motor centers, in mammals (right), dNG-ET^d^ and the hugely expanded iNG-ET^i^ (green) are intermixed and form mosaic circuits. While dNG-ET^d^ largely targets conserved structures in the forebrain and midbrain, iNG-ET^i^ massively amplifies, diversifies and *innovates* projection classes to brainstem and spinal cord. Moreover, iNG-ET^i^ expands projections in an area specific manner (extreme right) reflected in the expanded projections to action selection, diversification and execution centers from the motor areas; to sensory processing centers along the lemniscal pathways from the sensory areas; and to neuromodulatory, hypothalamic and other pallial centers from higher order areas of the isocortex. Abbreviations: dNG – direct neurogenesis; iNG - indirect neurogenesis; ET^d^ – direct neurogenesis generated extratelencephalic neuron; ET^i^ – indirect neurogenesis generated extratelencephalic neuron; MLR - mesencephalic locomotor region; RSN - reticulospinal neurons; TN - trigeminal nucleus; vlRN - ventro-lateral reticular nuc;eus; FMN - facial motor nucleus; LatRM - lateral rostral medulla; TSN - trigeminal sensory nucleus; SPN - spinal nucleus of the trigeminal; SpC – spinal cord.

A developmental and evolutionary (i.e. evo-devo) perspective is uniquely powerful in interpreting and understanding the stark differences between ET^d^ and ET^i^ projection patterns and provides deep insights into the basic organization of the cortical output system. In early vertebrates, the ancient pallium already evolved a blueprint of projections to downstream motor centers ^51^, with several cardinal cell types ^52^ and basic connectivity^53^, but the overall behavioral repertoire was limited and the involvement of pallium in motor control was rudimentary ^1,54^. Motor centers in the midbrain tectum (superior colliculus) ^55^, tegmentum and the well conserved BG in the forebrain ^56^, played key roles in motor control and behavior (**Figure 8**) ^57^. With the emergence of amniotes (reptiles and mammals), the “higher order” orchestration of motor control has gradually shifted to the cortical pallium ^58,59^, culminating in the mammalian isocortex. In parallel, with the evolution of limbs, hands and digits, command centers for specialized movements have expanded in the midbrain, BS, and SpC ^60–62^. The mammalian isocortex in particular, has seen a massive elaboration of efferent projections, which grant increasing monosynaptic control of downstream motor centers including the SpC ^8,31^. Importantly, this momentous “shift” of high-level motor control to the isocortex is mediated by the vastly expanded and diversified ET corticofugal projections and is correlated with the evolutionary expansion of iNG (**Figure 8**). While the amplification of neuronal production to expand cortical size is inherent to iNG, a major question is whether and how dNG and iNG differentially contribute to the diversification and organization of neuron types, in particular ET neurons that mediate cortical output channels driving behavior. Our basic premise is that the mouse dNG-ET^d^ reflects the mammalian instantiation of the evolutionarily more ancient system that has been conserved (albeit updated) across vertebrates, while iNG-ET^i^ represents the mammalian expansion and innovation.

Across different cortical areas and functional systems, ET^d^ and ET^i^ projections from the motor areas present strong interpretations for their functional distinction in the context of motor control systems ^1,63,64^. In particular, the rostral forelimb orofacial area (RFO) coordinates orofacial and forelimb actions in food manipulation, in part via ET neurons ^33,36,41,65,66^. Consistent with the subcortical projection differences between RFO ET^d^ (to mostly forebrain and midbrain regions) and ET^i^ (additionally to brainstem and spinal cord; **Figure 3G**), optogenetic activation reveals corresponding functional distinctions: while ET^d^ induced restricted movements of head and trunk, ET^i^ additionally elicited coordinated orofacial and forelimb movements that resemble feeding. iNG-ET^i^ thus not only amplifies ET^d^ cell numbers but also diversifies functional ET types, enabling coordinated multi-effector movements likely through recruitment of multiple action diversification/execution centers in the brainstem/spinal cord. It is plausible that ET^d^ might represent the conserved pan-vertebrate pathway that mainly targets the midbrain superior colliculus to regulate midline axis head-body orienting movements, and ET^i^ might encapsulate the mammalian isocortical innovation that expands to the BS and SpC to control and coordinate multi-effector forelimb-orofacial movement. Consistent with our finding, there is a direct correlation of improved forelimb dexterity with the expansion of the corticospinal neurons even between closely related rodent species^67^. In addition to brainstem and SpC, we further reveal strong ET^i^ over ET^d^ projection from motor areas (e.g. RFO, CFA, MOs) to all BG nuclei beyond the striatum, including GPe ^68^ and output nuclei ^69^ (**Figure 3G-I**) which may enhance cortical modulation of BG circuits during motor learning ^70^ and action selection^39,71^.

Sensory areas also show substantially expanded ET^i^ projections over ET^d^. The barrel cortex exhibits both strong CT (ET^d^) projection to FO and extensive PT (ET^i^) projections to HO, BS, and SpC. As a rodent-specific sensorimotor area ^72^, barrel cortex may exemplify the evolution of hybrid output pathways via iNG- ET^i^, consistent with isocortical expansion of sensory areas, ^3,73,74^. Notably, the primary visual cortex shows strong ET^i^ and minimal ET^d^ projections to the GABAergic prethalamic nuclei like vLGN, which is expanded in nocturnal mammals ^40^ and mediates innate visual behaviors such as fear responses^75,76^. Input to vLGN from visual cortical areas can suppress such responses ^77^, suggesting that iNG-ET^i^ diversification might enable the cortex to modulate instinctive behaviors through learning. iNG-ET^i^ expansion in sensory areas also targets the lemniscal pathways, such as principal trigeminal nucleus (barrelettes, ^78^ and VPM (barreloids, ^79^) targeted by barrel cortex, and the superior olive (sound localization, ^80^) and inferior colliculus (acoustic-motor functions^81^) targeted by auditory cortex. These ET^i^ projections may confer cortical modulation of ascending sensory processing at multiple levels.

In higher-order areas, the parietal cortex supports decision-making and movement planning ^41,82,83^, and shows expanded ET^i^ output to BG, BS, and SpC. PFC, the most recently evolved isocortical area that supports cognitive control ^84^, shows elaborate ET^i^ but few ET^d^ projections to pallial targets like BLA and hippocampus, structures critical for social behavior and memory ^85–88^, and to neuromodulatory centers like the VTA and dorsal raphe which can influence reward and behavioral states ^89^. PFC ET^i^ projections to the hypothalamus might confer cortical regulation of homeostatic and defensive behaviors^99,100^.

Altogether, our results reveal the systematic distinctions and mosaic organization of dNG-ET^d^ and iNG-ET^i^ projection streams across the isocortex. They suggest that the evolutionary expansion and diversification of iNG-ET^i^ may contribute to the expansion, diversification, and functional specialization of numerous isocortical areas through their massive and areal-differentiated corticofugal projections, which further shape the rest of the central nervous system architecture (**Figure 8**).

The area-specific ET projection patterns are likely shaped by the interplay of two broad classes of developmental mechanisms involving progressive as well as regressive processes. While genetic programs for targeted axon growth and guidance are well-suited to specify a species-invariant projection scaffold, regressive axon pruning allows for refinement of initially more exuberant branches, possibly in an activity- and utility-dependent manner to sculpt functional connections ^44^. Indeed, axon pruning has been implicated in the development of ET projection patterns ^46^. ET neurons across broad cortical areas initially all extend their axons distally to the spinal cord, then undergo massive axon pruning to sculpt their subcortical projection patterns and area specific distribution ^47,46,47^. A recent study examined axonal pruning of two broad ET subpopulations: ET_dist_ that predominate in the motor cortex and project distally to the pons, medulla and SpC, and ET_prox_ that predominate in visual cortex and project more proximally to pons and thalamus ^12,47^. Whereas subsets of ET_prox_ emerge from pruning of their initially distal-projection axons, others emerge through targeted growth. Here, we found that during the first postnatal week, ET^i^ neurons across the developing cortex undergo exuberant axon overgrowth to the spinal cord, followed by massive pruning to achieve areal specific distribution patterns in mature cortex. In sharp contrast, most developing ET^d^ across the cortex do not extend axons to the spinal cord, with the exception of a small subset largely restricted to motor areas. Notably, within the developing secondary motor and somatosensory areas, L6 in addition to L5 ET^i^ axons also project to the SpC, which are subsequently pruned (Figure 6C,D). Furthermore, beyond the isocortex, broadly defined Fezf2-expressing ET^i^ but not ET^d^ neurons in other cortical structures (e.g. hippocampus, BLA, entorhinal cortex, etc) also initially project to SpC and then undergo pruning.

Although we cannot rule out the possibility that a subset of ET^d^ neurons that project to more proximal targets also undergo pruning, our results indicate that exuberant axon growth followed by extensive pruning is a general and key feature that distinguish ET^i^ from ET^d^. It is plausible that substantial axon pruning is an effective mechanism to sculpt species-typic as well as individually customized ET projection patterns in each animal, in part through use- and activity-dependent mechanisms.

Beyond axon projection patterns, our reliable experimental access to dNG-ET^d^ and iNG-ET^i^ will enable multi-faceted analysis of these developmental genetic-defined subpopulations. Future studies will trace their synaptic input patterns, record and manipulate their activity during behavior, and reveal their gene expression profiles and regulatory programs. These studies will integrate the developmental genetic and systems neuroscience approaches toward understanding the diversity and organization of cortical neuron types in the context of vertebrate evolution.

Major unresolved questions include the evolutionary origin(s) of indirect neurogenesis and its massive expansion along the mammalian clade, including the underlying molecular genetic mechanisms that regulate the balance between dNG and iNG. Contrasting the previously held view of an amniote origin^28,90^, recent studies provide evidence for a more ancient evolution of iNG dating back to amphibians and even earlier to jawed vertebrates^26,27^. A previous study shows that the balance between dNG and iNG in the dorsal pallium is regulated by the activity levels of a conserved Robo/Slit and Notch/Delta-like 1 (Dll1) signaling across several amniote species^90^. Notably, comparative transcriptomics between shark and mouse revealed that RGs in both species exhibit high expression of Notch-family genes, while IOs express the Delta gene Dll1, suggesting that the Delta-Notch signaling axis constitutes an evolutionary toolkit for indirect neurogenesis that might have persisted for over 450 million years^26^. Intriguingly, mouse IPs exhibit an enhanced proliferative profile compared to their shark counterparts showing a stronger differentiation profile. This raises the possibility that evolutionary changes may have promoted the proliferative capacity in mammalian IPs, thereby contributing to the amplification and diversification of PN types that drive the expansion and complexification of the mammalian cortex.

## Acknowledgements

We thank Dr. Dhananjay Huilgol and Dr. Yi Li for helpful discussions. We are indebted to Leah Chestnut and Aditi Patel (LifeCanvas Technologies) for their excellent work on Lightsheet Microscopy Imaging. We thank Bao-Xia for her assistance with genotyping. Funding: NIH grant U19MH114823-01 (ZJH). NIH Director’s Pioneer Award 1DP1MH129954-01 (ZJH). International Postdoc Grant, Swedish Research Council grant no: 2021-00238 (SMS).

## Author Contributions

ZJH and SMS conceived of the study. SMS, ZJH, XA designed experiments. SMS and XA performed the experiments. YQ designed the *FezF2-TC* mouse line. SZ made the AAV viruses. SMS and XA analysed the data with inputs from HM. SMS and ZJH wrote the manuscript. ZJH supervised all aspects of the study and provided the funding.

## Declaration of interests

The authors declare no competing interests.

## Inclusion and diversity

We support inclusive, diverse, and equitable conduct of research.

## STAR Methods

### Key resources table

| REAGENT or RESOURCE | SOURCE | IDENTIFIER |
| --- | --- | --- |
| Antibodies |  |  |
| Chicken anti-GFP | Aves Labs | Cat# GFP-1020;<br>RRID:AB_10000240 |
| Rabbit anti-RFP | Rockland | Cat# 600-401-379;<br>RRID:AB_2209751 |
| EasyIndex | LifeCanvas Technologies | https://lifecanvastech.com/shop/easyindex/ |
| Nissl | ThermoFisher | Cat: N21483 |
| Bacterial and virus strains |  |  |
| AAVPHP.eB-TRE3G-EGFP | This study |  |
| AAVPHP.eB-CAG-FLEX-tdTomato | addgene | Catalog # 28306-PHPeB |
| AAV9-TRE3G-EGFP | This study | N/A |
| AAV9-CAG-FLEX-tdTomato | addgene | Catalog # 28306-AAV9 |
| AAVretro-TRE3G-mScarlet | This study | N/A |
| AAVretro-CAG-DIO-EGFP | addgene | Catalog # 50457-AAV9 |
| AAVretro-TRE3G-H2B-mScarlet | This study | N/A |
| AAVretro-CAG-DIO-H2B-EGFP | This study | N/A |
| AAVretro-CAG-EGFP | This study | N/A |
| Chemicals, peptides, and recombinant proteins |  |  |
| Tamoxifen | Sigma | T5648 |
| Goat serum | ThermoFisher | 10000C |
| Triton-X100 | Sigma | 9036-19-5 |
| Experimental models: Organisms/strains |  |  |
| FezF2-tTA-CreER | This study | N/A |
| Tbr2-2A-FlpO | JAX; Huilgol et al.<sup>17</sup> | RRID:IMSR_JAX:036297 |
| Fezf2-2A-CreER | JAX;Matho et al.<sup>32</sup> | RRID:IMSR_JAX:036296 |
| LSL-Flpo | JAX;He et al.<sup>91</sup> | RRID:IMSR_JAX:028584 |
| Intersection-Subtraction (IS2) | JAX;He et al.<sup>91</sup><br>Huilgol et al.<sup>17</sup> | RRID:IMSR_JAX:036760 |
| Ai32 | Madisen et al.<sup>48</sup> | RRID:IMSR_JAX:024109 |
| Software and algorithms |  |  |
| Matlab | MathWorks | https://www.mathworks.com/ |
| python | Python Software Foundation | https://www.python.org/ |
| CellPose | Pachitariu et al.<sup>92,93</sup> | https://cellpose.readthedocs.io/en/latest/ |
| DeepLabCut | Mathis et al.<sup>94</sup> | https://github.com/DeepLabCut/ |
| ZEN software | Zeiss | https://www.zeiss.com/microscopy/us/products/software/zeiss-zen.html |
| ImageJ/Fiji | Schindelin et al.<sup>95</sup> | https://imagej.net/software/fiji/ |

### Resource availability

#### Lead contact

Further information and requests for resources and reagents should be directed to and will be fulfilled by the lead contact, Z. Josh Huang.

#### Materials availability

Materials generated in this study are available on request to the lead contact. *FezF2-TC* mice generated in this study will be deposited to Jackson Laboratory.

#### Experimental model and study participant details

We have used adult mice (*Mus musculus*) in this study. Adult male and female mice bred onto a C57BL/6J background were used in the experiments. All mouse colonies were maintained in accordance with husbandry protocols approved by the IACUC and housed by gender in groups of 2–5 with access to food and water ad libitum and on a 12-hour light/dark cycle. The experimental procedures were approved by the Institutional Animal Care and Use Committee of Duke University and performed in accordance with the US National Institutes of Health (NIH) guidelines. All adult mouse histology experiments were performed on or after postnatal day P28-P40 animals. Postnatal experiments were performed on P1-P6 mice. We have used the following knock-in mice in this study: Driver lines include *FezF2-tTA-CreER* (this study);*Tbr2-2A-FlpO* (RRID:IMSR_JAX:036297); *Fezf2-2A-CreER* (RRID:IMSR_JAX:036296). Reporter lines include *Intersection-Subtraction (IS2)* (RRID: IMSR_JAX:036760) and *LSL-Flpo* (RRID:IMSR_JAX:028584). Mouse related experimental procedures were approved by the Institutional Animal Care and Use Committee (IACUC) of Duke University in accordance with NIH guidelines.

### Method details

#### Generation of *FezF2-tTA-CreER* knock-in mouse line

*FezF2-tTA-CreER* was generated by inserting a *frt-tTA-frt-CreER* cassette in frame before the STOP codon of *FezF2*. In *FezFTC;Tbr2-Flp* double*-*allele mice, dNG-derived ET^d^ express *tTA* while iNG-derived ET^i^ express *CreER*, as *tTA* is removed by *Tbr2-Flp* in IPs. Postnatal injection of *AAV-TRE-GFP* and *AAV-FLEX-RFP* in cortex differentially label ET^d^ and ET^i^ respectively. Targeting vectors were generated using a PCR-based cloning approach as described before ^17,32,91^.

#### Tamoxifen induction

Tamoxifen (T5648, Sigma) was prepared by dissolving the powder in corn oil (20 mg/ml) and constant magnetic stirring overnight at 37°C. A 100–200 mg/kg dose was administered by intraperitoneal injection at the appropriate age; If two doses, 100mg/kg doses were administered at P21 and P28. For experiments with early postnatal mice, it was administered intraperitoneally at P5-P14 from a diluted stock of 5mg/ml. For *FezF2TC;Tbr2Flp* mice with viral infections, tamoxifen induction was performed via intraperitoneal injections at a dose of 100 mg/kg with the first induction after one day following the viral infection and subsequently every 2-3 days, 2-3 times.

#### Immunohistochemistry

Adult mice were anaesthetized (using Isoflurane) and transcardially perfused with phosphate buffered saline (PBS) followed by 4% paraformaldehyde (PFA) in 0.1 M PBS. Post-fixation, brains were rinsed three times in PBS and sectioned at a 65-70μm thickness with a Leica VT1000S vibratome. Sections were treated with a blocking solution (10% normal goat serum and 0.2% Triton-X100 in 1X PBS) for 1h, then incubated overnight at 4°C with primary antibodies diluted in the blocking solution. Sections were washed three times in PBS and incubated for 2h at room temperature with corresponding secondary antibodies, Goat or Donkey Alexa Fluor 488, 594 or 647 (1:500, Life Technologies) and DAPI or Blue/Far-Red Neurotrace fluorescent Nissl stain to label nuclei (1:1000 in PBS, Life Technologies, 33342, Molecular Probes N21479). Sections were washed three times with PBS and dry-mounted on slides using Fluoromount-G (SouthernBiotech, 0100-01) mounting medium.

#### In situ hybridization

Hybridization chain reaction (HCR) in situ was performed as previously described^96^. *FezF2* probes were ordered from Molecular Instruments. Mouse brains were sliced into 50-µm-thick slices after paraformaldehyde (PFA) perfusion fixation and sucrose protection. HCR in situ was performed via the free-floating method in a 24-well plate. First, brain slices were exposed to a probe hybridization buffer with HCR Probe Set at 37::J°C for 24::Jhours. Brain slices were washed with a probe wash buffer, incubated with amplification buffer and amplified at 25::J°C for 24::Jhours. On day 3, brain slices were washed, counter-stained with DAPI/Nissl and mounted. *FezF2* (546::Jnm) probes were used to label *FezF2+* neurons.

#### Viral vectors

All viral vectors were produced in house. Viral Vectors used: *AAV9-CAG-FLEX-tdTomato*, *AAV9-TRE3G-EGFP, AAVPHP.eB-TRE3G-EGFP, AAVPHP.eB-CAG-FLEX-tdTomato, AAVretro-TRE3G-mScarlet, AAVretro-CAG-DIO-EGFP, AAVretro-TRE3G-H2B-mScarlet, AAVretro-CAG-DIO-H2B-EGFP, AAVretro-CAG-EGFP*. All viral vectors were aliquoted and stored at -80 °C until use.

#### Stereotaxic surgery

Mice, anesthetized with isoflurane (2-5 % at the beginning and 0.8-1.2 % for the rest of the surgical procedure), were positioned in a stereotaxic frame and their body temperature was maintained at 34-37 °C with a heating pad. Lidocaine (2%) was applied subcutaneously to the scalp prior to surgery. Ketoprofen (5 mg/kg) was administered intraperitonially (IP) as an analgesic before and after surgery. A vertical incision was made through the scalp and connective tissue to expose the dorsal surface of the skull. The skin was pushed aside, and the skull surface was cleared using saline. A digital mouse brain atlas, linked to the stereotaxic frame, guided the identification and targeting of different brain areas (Angle Two Stereotaxic System, Leica Biosystems). A small burr hole was drilled in the skull and brain surface was exposed. A pulled glass pipette, with a tip of 20-30 μm, containing the viral suspension was lowered into the brain. A 500-700 nl volume was delivered at a rate of 10-30 nl/min using a Picospritzer (General Valve Corp). The pipette remained in place for 5 min, to prevent backflow, prior to retraction. The incision was closed with Tissueglue (3M Vetbond) and 5/0 nylon suture thread (Ethilon Nylon Suture, Ethicon). The mice were kept warm on a heating pad during recovery. Coordinates used are as follows (all measured in mm from bregma except for medulla):

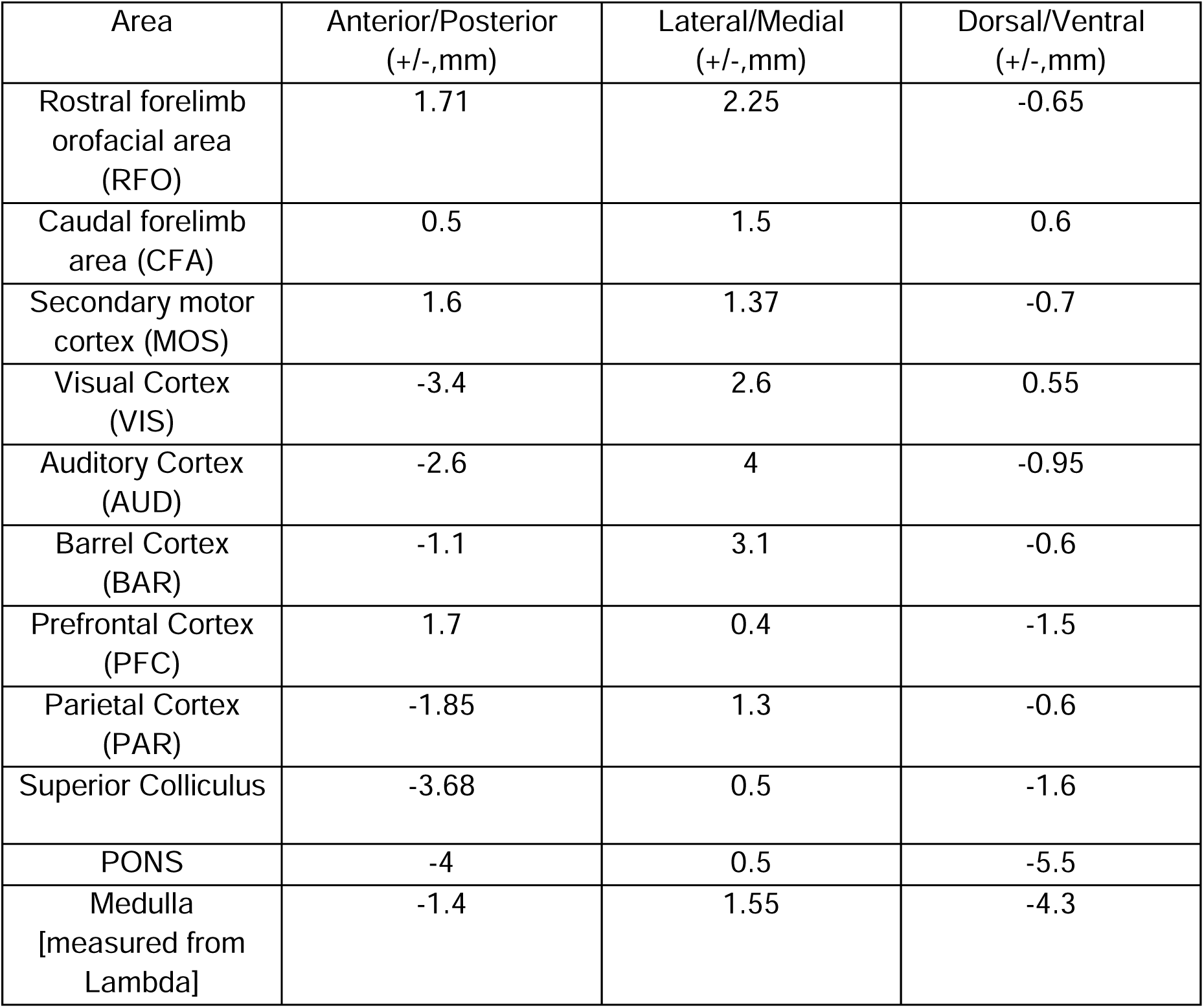

Injections into the spinal cord of adult mice were performed as described^97^. Specifically, a dorsal incision was made along the vertebral column at the cervical level. The overlying muscle was dissected to expose the cervical vertebral column, and a laminectomy was performed at C5-7 to expose the spinal cord. 0.5 μl of viral solution was injected at 0.4 mm lateral to midline and 0.5 mm and 1 mm depth using a glass micropipette. The dorsal muscle layers were sutured, and the skin was closed with sterile wound clips. For all anterograde and retrograde injections, a viral suspension containing a mixture of *AAV9-CAG-FLEX-tdTomato* and *AAV9-TRE3G-EGFP* or *AAVretro-CAG-DIO-EGFP* and *AAVretro-TRE3G-H2B-mScarlet* was used respectively, in the ratio 1:1. Injections into the cervical spinal cord of P1,P4 or P6 mice were performed using a Hamilton syringe (5 µl) with a mixture of *AAVretro-TRE3G-H2B-mScarlet* and *AAVretro-CAG-DIO-H2B-EGFP* (1:1, 2-2.5 µl) or just *AAVretro-CAG-EGFP* (1-2 µl) as described ^98^.

#### Light-Sheet Fluorescence Microscopy

For all anterograde and retrograde injections, post-fixed brains were processed for tissue-clearing, antibody staining and Lightsheet Scanning. A total of 36 brains were processed and imaged by LightSheet Microscopy (by LifeCanvas Technologies (LCT)).

#### Tissue Preservation, Clearing, Immunolabeling and Imaging

Paraformaldehyde-fixed samples were preserved using SHIELD reagents (LCT) using the manufacturer’s instructions ^99^. Samples were delipidated using LCT Clear+ delipidation reagents for 2 days. Following delipidation, samples were labeled using eFLASH^100^ technology which integrates stochastic electrotransport ^101^ and SWITCH ^102^, using a SmartBatch+ (or SmartLabel) device (LCT). For immunolabeling, the samples were incubated with 96 µg Chicken anti-GFP, 50 µg Goat anti-RFP, and 72 µg Rabbit NeuN (36 brains) following which the samples were incubated in 50% EasyIndex (RI = 1.52) overnight at 37°C followed by 1 d incubation in 100% EasyIndex for refractive index matching (RI = 1.52). After index matching the samples were imaged using a SmartSPIM axially swept light sheet microscope using a 3.6x objective (0.2 NA) with 4 um z-step, and 1.8 um x-y pixel size.

#### Atlas Registration (NeuN)

Samples were registered to the Allen Brain Atlas^103^ using an automated process (LCT). A NeuN channel for each brain was registered to an average NeuN atlas (generated by using previously registered samples). Registration was performed using successive rigid, affine, and b-spline warping algorithms (SimpleElastix: https://simpleelastix.github.io/ ).

#### Cell Detection

Automated cell detection was performed by using a custom convolutional neural network created with the Tensorflow python package (Google, LCT). The cell detection was performed by two networks in sequence. First, a fully-convolutional detection network ^104^ based on a U-Net architecture ^105^ was used to find possible positive locations. Second, a convolutional network using a ResNet architecture ^106^ was used to classify each location as positive or negative. Using the previously calculated Atlas Registration, each cell location was projected onto the Allen Brain Atlas in order to count the number of cells for each atlas-defined region.

#### Image Segmentation and Thresholding

Process-specific signals were identified by applying an absolute intensity threshold to each voxel within the volumetric image. Voxels exhibiting fluorescence intensity greater than the predetermined threshold were classified as “positive.”

#### Atlas Registration (Anterograde projections)

The pixels from the resulting segmentation mask were spatially transformed (warped) to the Allen Brain Atlas to determine which region each positive pixel belongs to.

#### Computation of Fraction Occupation and Percent Occupation

For each anatomical region defined by the registered atlas, the number of positive voxels (i.e., those exceeding the intensity threshold) was summed. This count was then divided by the total voxel count of the same region to yield the fraction occupation metric:

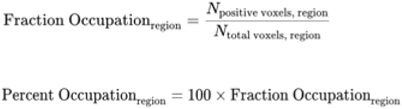

#### Coronal Video Reconstruction

Whole Brains were imaged by LightSheet Microscopy horizontally at a resolution of 1.8, 1.8, and 4 um for the medial-lateral, anterior-posterior, and dorsal-ventral axes respectively. The acquired whole-brain image data was resampled to generate a series of raw coronal images with a resolution of 1.8 and 4 um for the medial-lateral and dorsal-ventral axes respectively. The dorsal-ventral axis was downsampled to a resolution of 54 um. To make the video of coronal images, we further downsampled each raw coronal image five times. The display range of each color channel was carefully adjusted linearly to reveal axons and cell bodies of anterograde and retrograde tracing data respectively.

#### Confocal Microscopy

Confocal photomicrographs were obtained using a Zeiss Laser scanning microscope (LSM 900, in house). Tile-scans were taken with a 5x objective while Z-stacks of optical sections were obtained using either a 10x or a 20x objective. Projection images and illustrations were processed/prepared by Zeiss ZEN software, ImageJ and Adobe Illustrator CC. Various sections of interest from a total of 40 mouse brains were analyzed using confocal microscopy.

#### Optogenetic activation of ET^all^, ET^d^ and ET^i^

For optogenetic activation, ChRmine was expressed in ET^d^ on the ipsilateral side of RFO using viral injections of AAV8-TRE3G-ChRmine-oScarlet and in ET^i^ using AAV8-nEF-FLEX-ChRmine-oScarlet on the contralateral side of RFO in the same *FezF2TC;Tbr2Flp* mice. For optogenetic activation of ET^all^, *FezF2CreER* mice were crossed with the *Ai32* reporter mice to express channelrhodopsin in all *FezF2^+^* neurons. Tapered optical fibers (diameter 200 μm; NA, 0.37; taper 1 mm) were implanted bilaterally in the RFO. The optical fibers were implanted at a depth 650 μm from the cortical surface. To fix the implants to the skull, a silicone adhesive (Kwik-Sil, WPI) was applied to cover the hole, followed by a layer of dental cement (C&B Metabond, Parkell), black instant adhesive (Loctite 426), and dental cement (Ortho-Jet, Lang Dental). A titanium head bar was fixed to the skull near Lambda using dental cement.

During head-fixed activation, two cameras (FL3-U3-13S2C-CS, FLIR) were placed in front of the mice. Two LED light lamps next to the two cameras provided illumination. A fiber coupled laser (5-ms pulses, 10-20 mW; λ = 473/630 nm) was used to apply stimulation at 10, 20, 30, and 50 Hz and constantly for 0.5 s. For free-moving activation, mice with optical fibers implanted in the RFO were placed into an acrylic activity box (14 cm × 14 cm × 16.5 cm, L × W × H). A 473-nm laser (5-ms pulses, 10 mW) coupled to a rotary joint (RJPFL2, Thorlabs) was used to apply stimulation at 10, 20, 30, and 50 Hz and constantly for 0.5 s. One camera (FL3-U3-13S2C-CS, FLIR) was used to take video recordings at a frame rate of 100 Hz from the bottom of the activity box. A LED light lamp adjacent to the camera provided illumination.

### Data Analysis

#### Heatmap Visualization of Projection Patterns

For anterograde projections, fraction occupation percentage data (obtained from the LightSheet analysis pipeline, LCT) were extracted for selected ipsilateral and contralateral brain regions for each injection site. For each injection site, the downstream areas selected across the brain were based on comparisons of equivalent injections (based on coordinates) in the isocortex from the Allen Mouse Brain Connectivity dataset (https://community.brain-map.org/t/api-allen-brain-connectivity/2988). The downstream areas selected for each injection site are listed in Table S1. The data were then normalized, and the mean GFP and RFP values were calculated for each region. For global normalization, Min-Max scaling was applied to the combined GFP and RFP data across both ipsilateral and contralateral regions. The final normalized data were saved for further analysis. Normalized mean fraction occupation for ET^d^ and ET^i^ were visualized as heatmaps, for both ipsilateral and contralateral projections across different selected regions. Region names were aligned along the y-axis, and color intensity scaled to reflect normalized fraction occupation.

#### Linear Regression and Visualization

ET^d^ and ET^i^ fraction occupations for each injection site from target structures of interest (eg: thalamus, midbrain, PONS, medulla were extracted for both ipsilateral and contralateral regions. Structures selected for each of the target structures considered for each injection site are listed in Table S1. For each target structure, linear regression (polyfit) was performed between ET^d^ and ET^i^ fraction occupation values. Regression lines were plotted alongside scatter plots of individual data points representing ipsilateral and contralateral areas within each region group. Anatomical area labels were overlaid on the plots using text annotations. ET^d^ and ET^i^ fraction occupation values for two region groups compared were normalized using z-score transformation. The z-score for each data point was calculated using the formula:

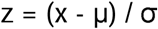

where x is the data point, μ is the global mean of the two region groups, and σ is the global standard deviation of two region groups. Percent occupation differences (Δ = ET^d^ - ET^i^) were computed for each target structure. Statistical tests were performed within-group and visualized through boxplots with median, interquartile range, and individual data points along with p-value annotations to indicate statistical significance. To further assess whether the percent occupation differences (Δ = ET^d^ - ET^i^) were uniform across different target structures from an injection site, a multicomparisons test followed by post-hoc comparisons were performed. Post-hoc comparisons were done between different target structures and the thalamus. Since the thalamus in general received significant ET^d^ and ET^i^ projections, it was used as a control for comparison with other target structures.

#### Cell Count Quantification Using Cellpose

Cell body segmentation and quantification for confocal images were performed using the Cellpose algorithm ^92^, a deep learning-based tool for cellular segmentation. Images were preprocessed for contrast enhancement and background subtraction before input into Cellpose. The “cyto” model was used with default parameters, and the estimated cell diameter was manually specified or automatically determined, depending on image resolution.

Batch segmentation was conducted via the Cellpose GUI or command-line interface, and outputs were visually inspected for accuracy. Resulting masks were used to quantify total cell counts per region or image.

#### Correlation Analysis

Correlation analysis was performed on either projection fraction occupation values or cell counts across brain regions, on brains scanned by Lightsheet Microscopy. For each subject, direct and indirect data from both hemispheres were combined. Correlation matrices were computed using Pearson correlation coefficients and visualized as heatmaps to compare projection/cell count similarities across individual mice.

#### Video analysis for optogenetic activation

Videos of behavior from head-fixed activation were analyzed with DeepLabCut^94^. The two cameras were calibrated using the Camera Calibrator App in MATLAB. For DeepLabCut training, 750 images were used from the frontal video recordings to track the movements of the jaw and hands. The tracking results were validated manually and errors were corrected accordingly in MATLAB. Trials in which mice made spontaneous movements before stimulation onset (within 0.5 s) were excluded from the analyses, based on either manual examinations or setting threshold (3 × SD from the mean) on average speed distributions of all trials. For videos obtained from free-moving activation, the tracking of different body parts was performed using DeepLabCut. The network was trained with 500 images. Seven body parts (left and right hands and feet, nose tip, jaw, and tail base) were labeled in the images. The behavioral videos and tracking results were visualized and analyzed in a custom-written MATLAB app. Tracking errors were corrected using the app.

#### Statistical analysis

For statistical comparisons, normality of the dataset was assessed using either the Shapiro-Wilk test or the Kolmogorov-Smirnov test. When data were not normally distributed, non-parametric statistical tests were used. For multiple comparisons, normality was assessed using the Lilliefors test followed by a nonparametric Kruskal-Wallis test. Post-hoc comparisons of different target regions with thalamus were performed by pairwise Wilcoxon rank-sum tests or Kolmogorov-Smirnov (KS) tests. To control for multiple comparisons while preserving statistical power, all post hoc p-values were corrected using the Benjamini-Hochberg false discovery rate (FDR) method. Significance levels used in the analyses and figures were as follows: *P < 0.05, **P < 0.01, ***P < 0.005, ****P < 0.001, *****P < 0.0001, and n.s. (not significant). Data are presented as median with the 25th–75th percentiles (IQR), or as mean ± s.d., unless otherwise indicated. All statistics were implemented in MATLAB.

## Data availability/Code availability

The data that support the findings of this study are available from the lead contact upon reasonable request. Custom-written scripts used in this study are available from: https://github.com/ShreyasDuke/ETd-ETi-Code.git. Anterograde projection and cell distribution video datasets can be viewed at: https://figshare.com/s/4aa9d712da9756bc7551.

## Supplemental Figure Legends

**Figure S1.**
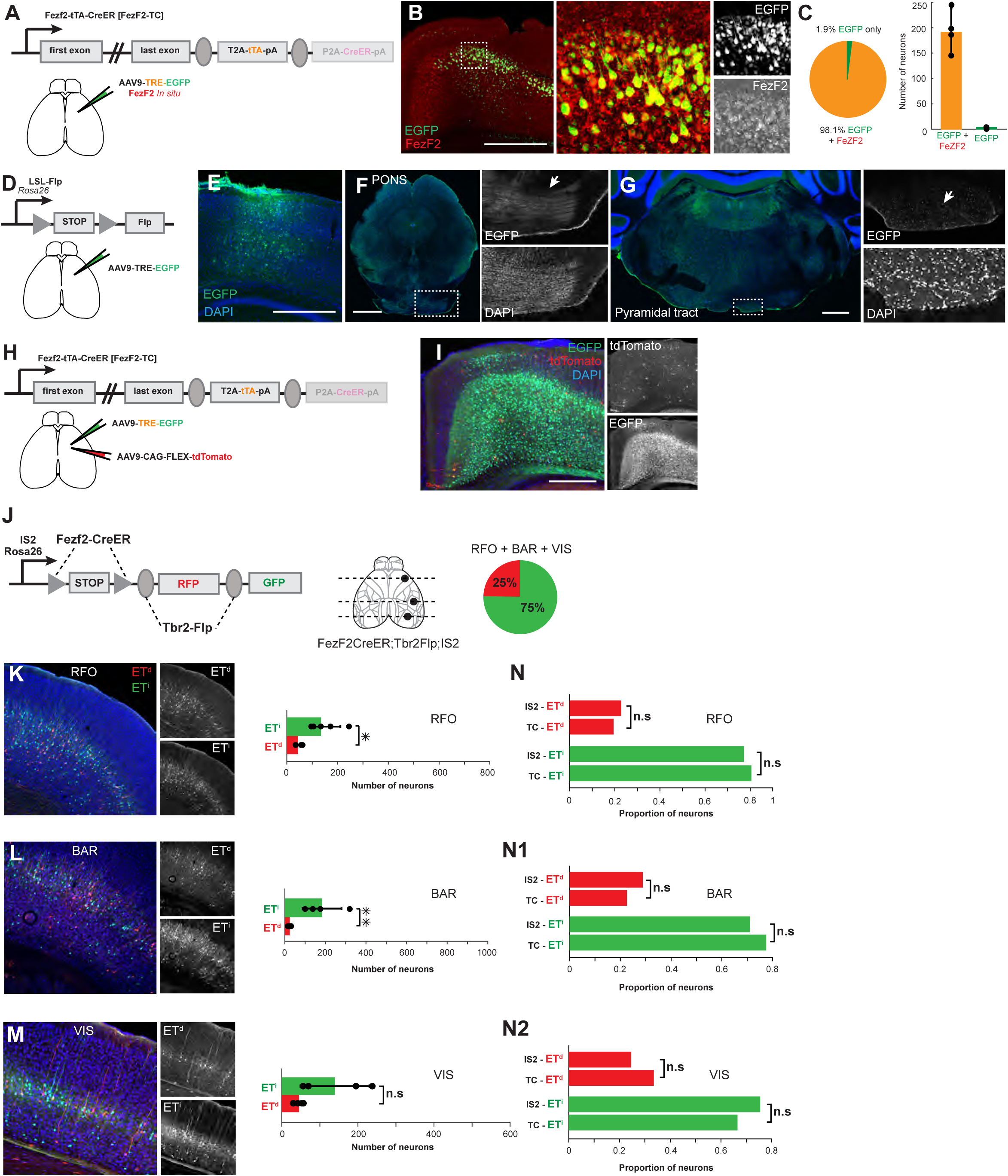
Validation of the *FezFTC;Tbr2Flp* mice for viral access to capture ET^d^ and ET^i^, Related to Figure 1. (A) Schematic of strategy for validating the *FezF2-TC* line. Without the*Tbr2-Flp* allele *FezF2-TC* drives *tTA* expression in all *FezF2+* neurons, which can be labelled by injection of *AAV-TRE-EGFP*. (B-C) The vast majority of the EGFP-labelled neurons in the deep layers overlap with *in situ* detection of *FezF2* mRNA (red) verifying that they are *FezF2+* (yellow). (D-G) Strategy and result of checking for non-specific labelling for *AAV-TRE-EGFP*. Injection of *AAV9-TRE-EGFP* in *LSL-Flp* mice, which do not express tTA (D), did not result in prominent GFP expression. Faint GFP signals at the injection site (E) resulted from low level leakage of *TRE-EGFP* but did not label any corticofugal axons in pons (F) and medulla (G) even after antibody amplification of EGFP signal. (H-I) Strategy and result for checking non-specific labelling for *AAV-CAG-FLEX-tdTomato*. We inject *AAV9-CAG-FLEX-tdTomato* together with *AAV9-TRE-EGFP* in *FezF2-TC* mice as in (A). All *FezF2+* neurons are labelled with EGFP at the injection site. There is negligible labelling of *tdTomato, as* there is no *Cre* expression in the *FezF2-TC*, indicating that there is almost no leaky expression. (J) (left) Schematic of a previous strategy wherein dNG and iNG can be simultaneously visualized by a genetic fate-mapping scheme using the IS2 reporter with *FezF2-CreER* (RG) and *Tbr2-Flp* (IP) drivers: dNG-ET^d^ (*Fezf2^+^*/*Tbr2-*) is labeled by RFP through “Cre-NOT-Flp” subtraction; iNG-ET^i^ (*FezF2^+^*/*Tbr2^+^*) is labeled by EGFP through “Cre-AND-Flp” intersection (see^17^). (right) Overall proportions of ET^d^ and ET^i^ neurons from RFO, BAR and VIS areas was 1:3. (K-M) (left) We compared sections from RFO (K), BAR (L) and VIS (M) corresponding to the sections in Figure 1C1-C3. (right) The number of ET^i^ neurons were significantly more than ET^d^ for RFO and BAR. (N-N2) Comparing proportions of dNG-ET^d^ and iNG-ET^i^ labelled from the new and old strategies: viral targeting in *FezFTC;Tbr2Flp* in Figure 1A and *FezFTC;Tbr2Flp;IS2* in (J), showed no significant differences showing both strategies are comparable and consistent for (N) RFO, (N1) BAR and (N2) VIS. Abbreviations: RFO – rostral forelimb orofacial area; BAR – barrel cortex; VIS – visual cortex; Error bars indicate SD. Scatter points indicate individual data. Statistical tests used: two-sample t-tests (p-values: *p < 0.05, **p < 0.01, n.s = not significant).Scalebars in (B,E,I) 500 µm (F,G) 1 mm.

**Figure S2.**
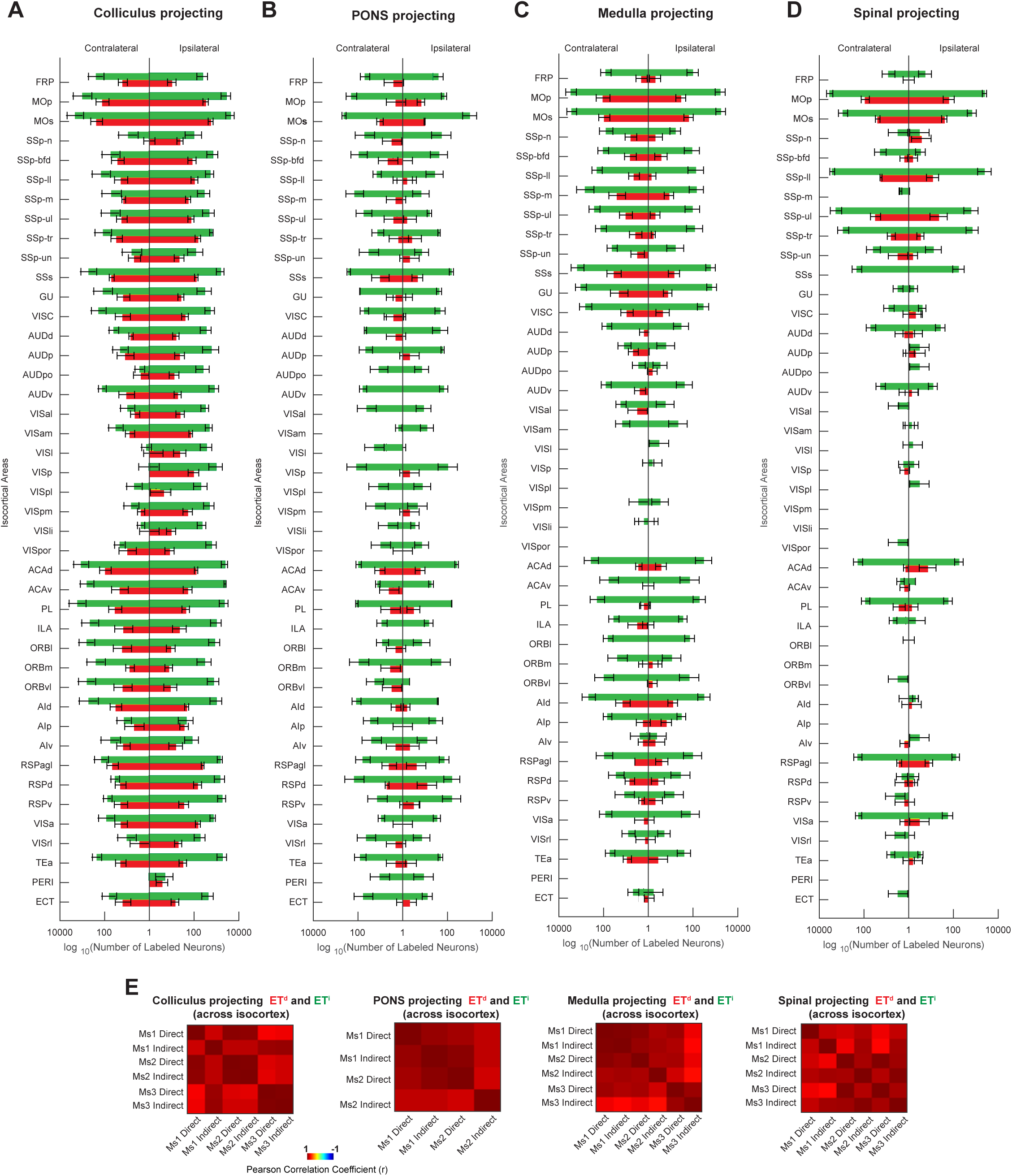
ET^d^ and ET^i^ collicular-, pontine-, medulla- and spinal- projecting neurons. Related to Figure 2. (A-D) Bar plots of retrogradely labelled ET^d^ (red) and ET^i^ (green) neuron numbers across different areas of the isocortex from collicular (A), pontine (B), medulla (C), and cervical spinal cord (D) injections. X-axis is shown in a Log_10_ scale. (E) Correlation matrix heatmap showing pairwise Pearson correlations between number of retrogradely labelled ET^d^and ET^i^ neurons across all areas of the isocortex from 2-3 mice each from collicular, pontine, medulla, and cervical spinal cord injections. Abbreviations for all isocortical regions in (A-D) are listed in Table S1. Error-bars represent SEM.

**Figure S3.**
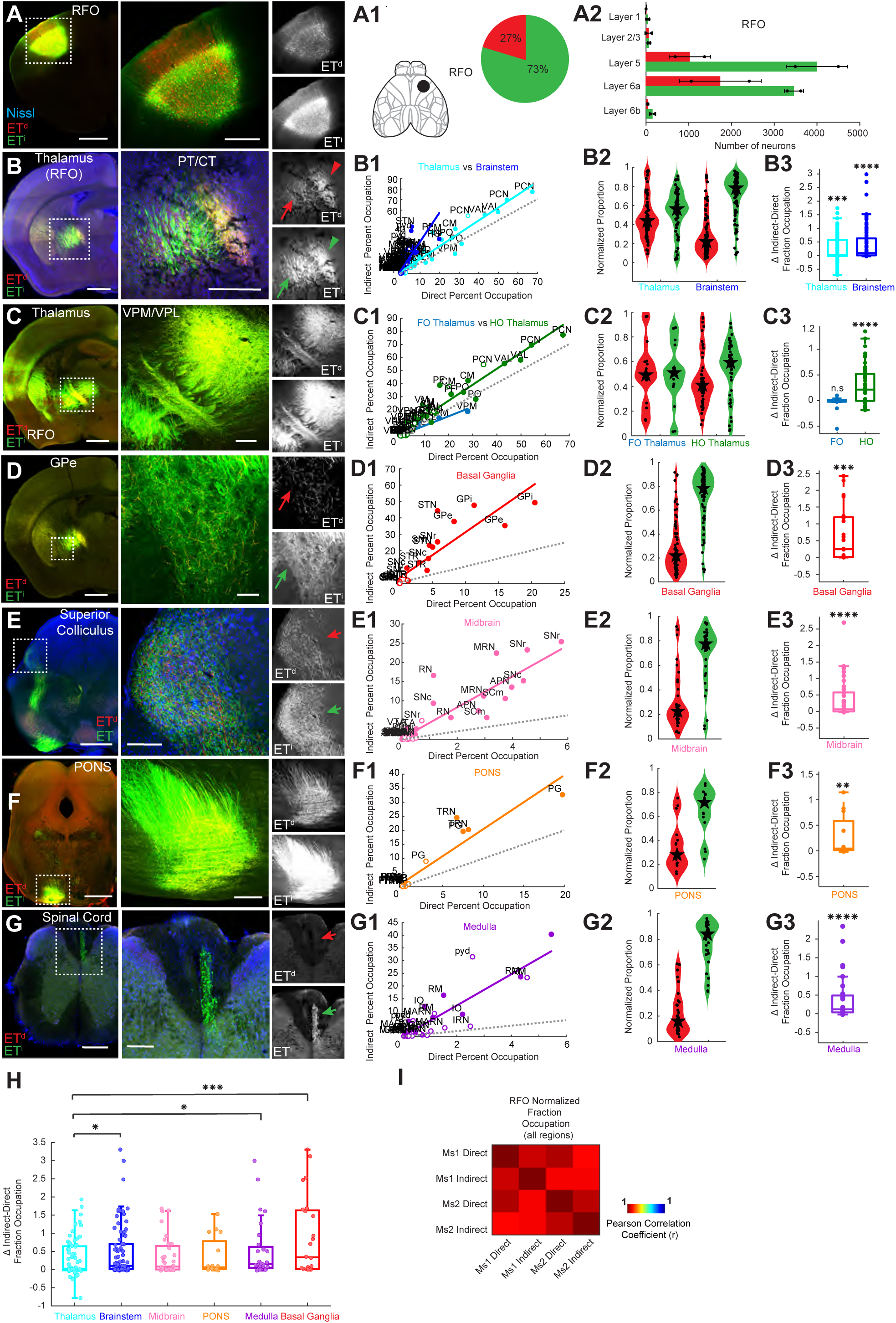
Predominant ET^i^ over ET^d^ subcortical projections from RFO, Related to Figure 3. (A) AAV injection in the RFO labeled ET^d^ (red) and ET^i^ (green). (A1) Left, location of the injection site shown on a schematic of the dorsal cortex. Right, pie chart of proportion of ET^d^ and ET^i^ neurons labelled at the injection site. (A2) Bar plots of layer-wise counts of labelled ET^d^ and ET^i^ neurons in the injection site (n = 2 mice). Labelling is evident in deep layers including layer 6. *FezF2* is known to recruit *Tle4* to repress PT genes, and their co-expression in layer 6^80^ likely explains their labeling. (B) PT and CT fiber bundles with ET^d^ and ET^i^ projections from RFO. Notice both ET^d^ and ET^i^ projections in CT (arrowhead) while largely ET^i^ in PT (arrow). (C) Dense ET^d^ and ET^i^ projections to the thalamus, including VPM/VPL. (D) Dense ET^i^ but weak ET^d^ projections to the GPe. (E) Stronger ET^i^ over ET^d^ projections to the superior colliculus. (F) Strong ET^i^ but weak ET^d^ projections to the PONS. (G) Strong ET^i^ but nearly no ET^d^ projections to the spinal cord. (B1-G1) Scatter plots with linear regression of normalized fraction occupations of RFO ET^d^ and ET^i^ projections to target regions (B1) thalamus and brainstem, (C1) first order (FO) and higher order (HO) thalamus, (D1) basal ganglia, (E1) midbrain, (F1) PONS, (G1) medulla. (B2-G2) Violin plots showing normalized percent occupation for RFO ET^d^ and ET^i^ projections with the star indicating the median and black circles showing individual values for subregions within target regions of (B2) thalamus and brainstem (C2) first order (FO) and higher order (HO) thalamus (D2) basal ganglia (E2) midbrain (F2) PONS (G2) medulla. (B3-G3) Box plots of Z-score normalized percent occupation differences between ET^i^ and ET^d^ projections from RFO to (B3) thalamus and brainstem, (C3) first order (FO) and higher order (HO) thalamus, (D3) basal ganglia, (E3) midbrain, (F3) PONS, (G3) medulla. p-values reflect significant within region differences. (H) Box plots of Z-score normalized percent occupation differences between ET^i^ and ET^d^ projections from RFO for brainstem, midbrain, PONS, medulla and BG with post-hoc comparisons to thalamus, following a multiple comparisons test (p < 0.05) to assess statistical differences among different projection target regions. The brainstem, medulla and BG regions showed significantly more iNG-ET^i^ projections than thalamus. (I) Correlation matrix heatmap showing pairwise Pearson correlations between ET^d^ and ET^i^ normalized fraction occupation across all areas for projections from RFO across two mice used for LightSheet Imaging. Abbreviations: RFO – rostral forelimb orofacial area; FO – first order thalamic nuclei; HO- higher order thalamic nuclei; VPM – ventral posterior medial nucleus; VPL – ventral posterior lateral nucleus; PO – posterior thalamic nucleus GPe- Globus pallidus externa; PONS – pontine nuclei; BG – basal ganglia. Subregions for each target region are listed in Table S1. Scatter points in scatter plots, violin plots and box plots represent individual subregion percent occupation (normalized or raw) values, with filled markers for ipsilateral and open markers for contralateral data. Regression lines indicate overall correlation trends. Dotted grey line indicates the line of equality. Interquartile range in box plots is 25th–75th percentiles. Statistical tests used one-sample t-test, Wilcoxon signed-rank test, Kruskal–Wallis test, Wilcoxon rank-sum test and Kolmogorov-Smirnov test (p-values: *p < 0.05, **p < 0.01, ***p < 0.001, ****p < 0.0001, n.s = not significant) Scalebars in (A-F) 1 mm; (G, A right, B right) 500 µm; (C right, E right, F right) 200 µm; (D right, G right) 100 µm.

**Figure S4.**
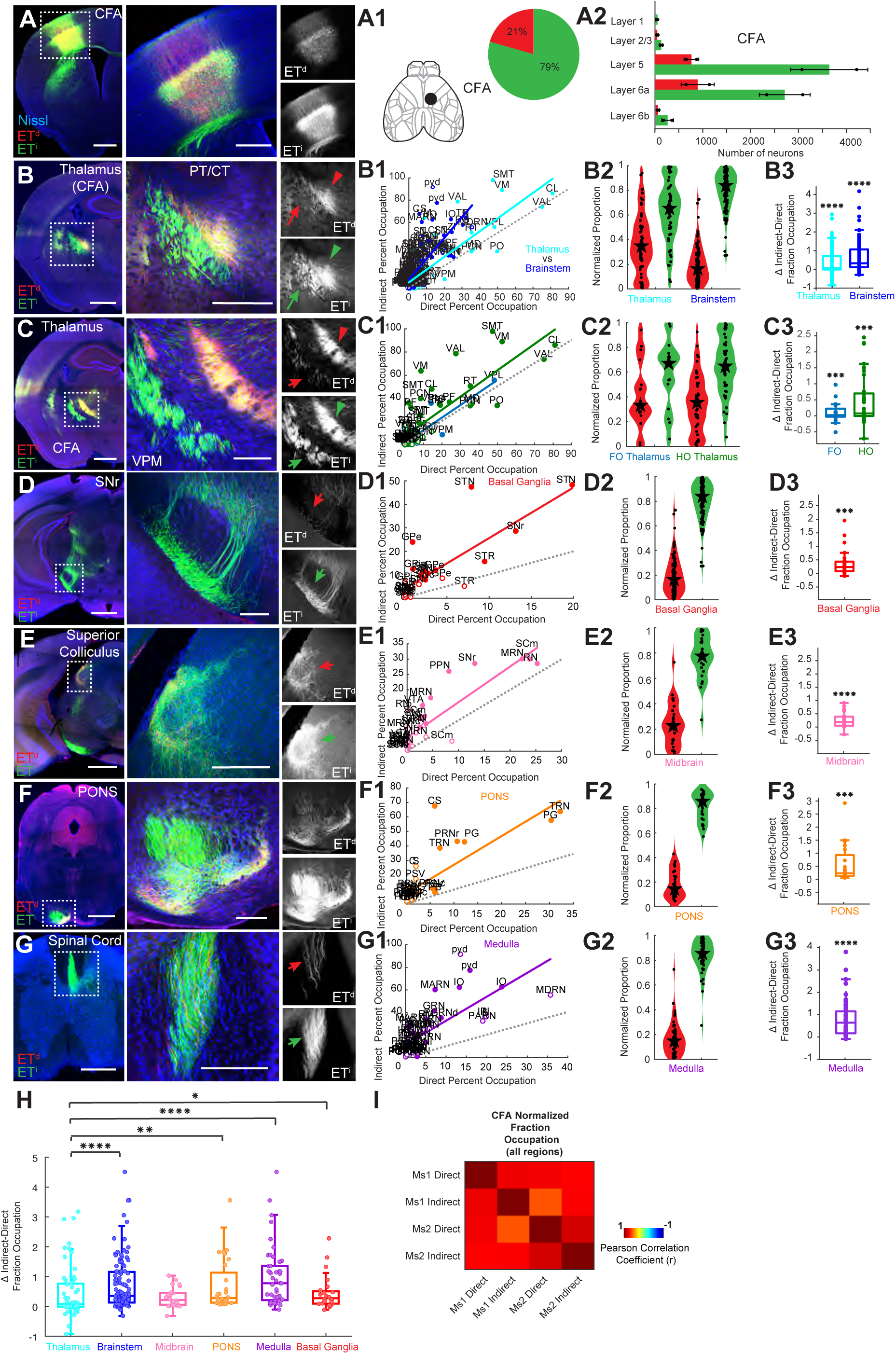
Predominant ET^i^ over ET^d^ subcortical projections from CFA, Related to Figure 3. (A) AAV injection in the CFA labelling ET^d^ (red) and ET^i^ (green). (A1) (left) Location of the injection site shown on a schematic of dorsal cortex. (right) Pie chart of proportion of ET^d^and ET^i^ neurons labelled at the injection site. (A2) Bar plots of layer-wise counts of labelled ET^d^ and ET^i^ neurons in the injection site (n = 2 mice). (B) PT and CT fiber bundles with ET^d^ and ET^i^ projections from CFA. Notice both ET^d^ and ET^i^ projections in CT (arrowhead) while largely ET^i^ in PT (arrow). (C) Dense ET^d^ and ET^i^ projections to the thalamus, including VPM. (D) Dense ET^i^ but weak ET^d^ projections to the SNr. (E) Stronger ET^i^ over ET^d^ projections to the superior colliculus. (F) Strong ET^i^ but weak ET^d^ projections to the PONS. (G) Strong ET^i^ but nearly no ET^d^ projections to the spinal cord. (B1-G1) Scatter plots with linear regression of normalized fraction occupations of CFA ET^d^ and ET^i^ projections to target regions (B1) thalamus and brainstem, (C1) first order (FO) and higher order (HO) thalamus, (D1) basal ganglia, (E1) midbrain, (F1) PONS, (G1) medulla. (B2-G2) Violin plots showing normalized percent occupation for CFA ET^d^ and ET^i^ projections with the Star indicating the median and black circles showing individual values for subregions within target regions of (B2) thalamus and brainstem (C2) first order (FO) and higher order (HO) thalamus (D2) basal ganglia (E2) midbrain (F2) PONS (G2) medulla. (B3-G3) Box plots of Z-score normalized percent occupation differences between ET^i^ and ET^d^ projections from CFA to (B3) thalamus and brainstem, (C3) first order (FO) and higher order (HO) thalamus, (D3) basal ganglia, (E3) midbrain, (F3) PONS, (G3) medulla. p-values reflect significant within region differences. (H) Box plots of Z-score normalized percent occupation differences between ET^i^ and ET^d^ projections from CFA for brainstem, midbrain, PONS, medulla and BG with post-hoc comparisons to thalamus, following a multiple comparisons test (p < 0.0001) to assess statistical differences among different projection target regions. The brainstem, PONS, medulla and BG regions showed significantly more iNG-ET^i^ projections than thalamus. (I) Correlation matrix heatmap showing pairwise Pearson correlations between ET^d^ and ET^i^ normalized fraction occupation across all areas for projections from CFA across two mice used for LightSheet Imaging. Abbreviations: CFA – caudal forelimb area; FO – first order thalamic nuclei; HO- higher order thalamic nuclei; VPM – ventral posterior medial nucleus; PO – posterior thalamic nucleus SNr – substantia nigra *pars reticulata*; PONS – pontine nuclei; BG – basal ganglia. Subregions for each target region are listed in Table S1. Scatter points in scatter plots, violin plots and box plots represent individual subregion percent occupation (normalized or raw) values, with filled markers for ipsilateral and open markers for contralateral data. Regression lines indicate overall correlation trends. Dotted grey line indicates the line of equality. Interquartile range in box plots is 25th–75th percentiles. Statistical tests used one-sample t-test, Wilcoxon signed-rank test, Kruskal–Wallis test, Wilcoxon rank-sum test and Kolmogorov-Smirnov test (p-values: *p < 0.05, **p < 0.01, ***p < 0.001, ****p < 0.0001, n.s = not significant) Scalebars in (A-F) 1 mm; (G, A-B right) 500 µm; (C right, E right, G right) 250 µm; (D right, F right) 200 µm.

**Figure S5.**
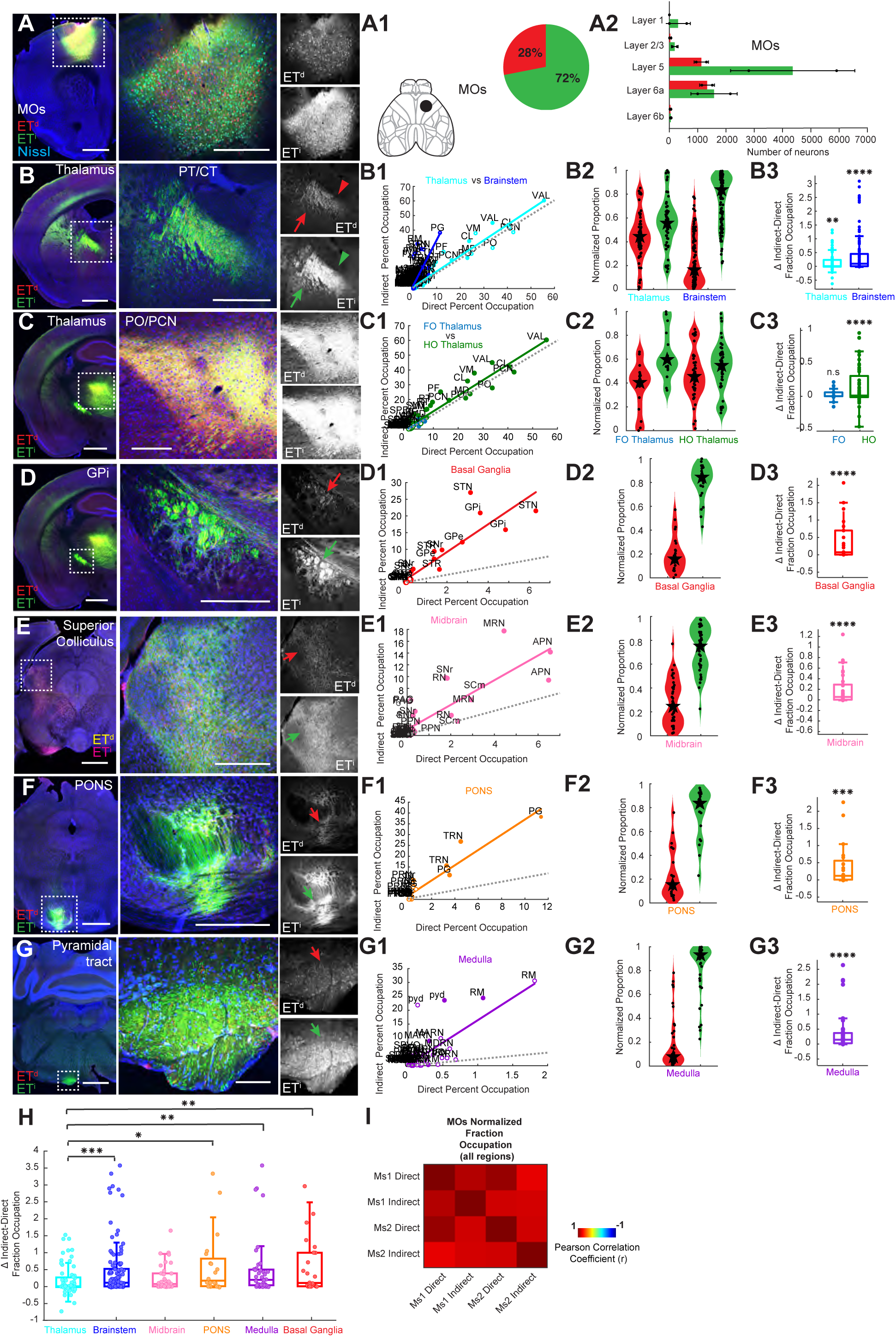
Predominant ET^i^ over ET^d^ subcortical projections from MOs, Related to Figure 3. (A) AAV injection in the MOs labeled ET^d^ (red) and ET^i^ (green). (A1) Left, location of the injection site shown on a schematic of dorsal cortex. Right, pie chart of proportion of ET^d^ and ET^i^ neurons labelled at the injection site. (A2) Bar plots of layer-wise counts of labelled ET^d^ and ET^i^ neurons in the injection site (n = 2 mice). (B) PT and CT fiber bundles with ET^d^ and ET^i^ projections from MOs. Notice both ET^d^ and ET^i^ projections in CT (arrowhead) while largely ET^i^ in PT (arrow). (C) Dense ET^d^ and ET^i^ projections to the thalamus, including PO/PCN. (D) Dense ET^i^ but weak ET^d^ projections to the GPi. (E) Stronger ET^i^ over ET^d^ projections to the superior colliculus. (F) Strong ET^i^ but weak ET^d^ projections to the PONS. (G) Strong ET^i^ but nearly no ET^d^ projections to the spinal cord. (B1-G1) Scatter plots with linear regression of normalized fraction occupations of MOs ET^d^ and ET^i^ projections to target regions (B1) thalamus and brainstem, (C1) first order (FO) and higher order (HO) thalamus, (D1) basal ganglia, (E1) midbrain, (F1) PONS, (G1) medulla. (B2-G2) Violin plots showing normalized percent occupation for MOs ET^d^ and ET^i^ projections with the Star indicating the median and black circles showing individual values for subregions within target regions of (B2) thalamus and brainstem (C2) first order (FO) and higher order (HO) thalamus (D2) basal ganglia (E2) midbrain (F2) PONS (G2) medulla. (B3-G3) Box plots of Z-score normalized percent occupation differences between ET^i^ and ET^d^ projections from MOs to (B3) thalamus and brainstem, (C3) first order (FO) and higher order (HO) thalamus, (D3) basal ganglia, (E3) midbrain, (F3) PONS, (G3) medulla. p-values reflect significant within region differences. (H) Box plots of Z-score normalized percent occupation differences between ET^i^ and ET^d^ projections from MOs for brainstem, midbrain, PONS, medulla and BG with post-hoc comparisons to thalamus, following a multiple comparisons test (p < 0.01) to assess statistical differences among different projection target regions. The brainstem, PONS, medulla and BG regions showed significantly more iNG-ET^i^ projections than thalamus. (I) Correlation matrix heatmap showing pairwise Pearson correlations between ET^d^ and ET^i^ normalized fraction occupation across all areas for projections from MOs across two mice used for LightSheet Imaging. Abbreviations: MOs – secondary motor area; FO – first order thalamic nuclei; HO- higher order thalamic nuclei; VPM – ventral posterior medial nucleus; VPL – ventral posterior lateral nucleus; PO – posterior thalamic nucleus; GPi – globus pallidus interna; PONS – pontine nuclei; BG – basal ganglia. Subregions for each target region are listed in Table S1. Scatter points in scatter plots, violin plots and box plots represent individual subregion percent occupation (normalized or raw) values, with filled markers for ipsilateral and open markers for contralateral data. Regression lines indicate overall correlation trends. Dotted grey line indicates the line of equality. Interquartile range in box plots is 25th–75th percentiles. Statistical tests used: one-sample t-test, Wilcoxon signed-rank test, Kruskal–Wallis test, Wilcoxon rank-sum test and Kolmogorov-Smirnov test (p-values: *p < 0.05, **p < 0.01, ***p < 0.001, ****p < 0.0001, n.s = not significant) Scalebars in (A-G) 1 mm; (A-B right, D-F right) 500 µm; (C right) 250 µm; (G right) 100 µm.

**Figure S6.**
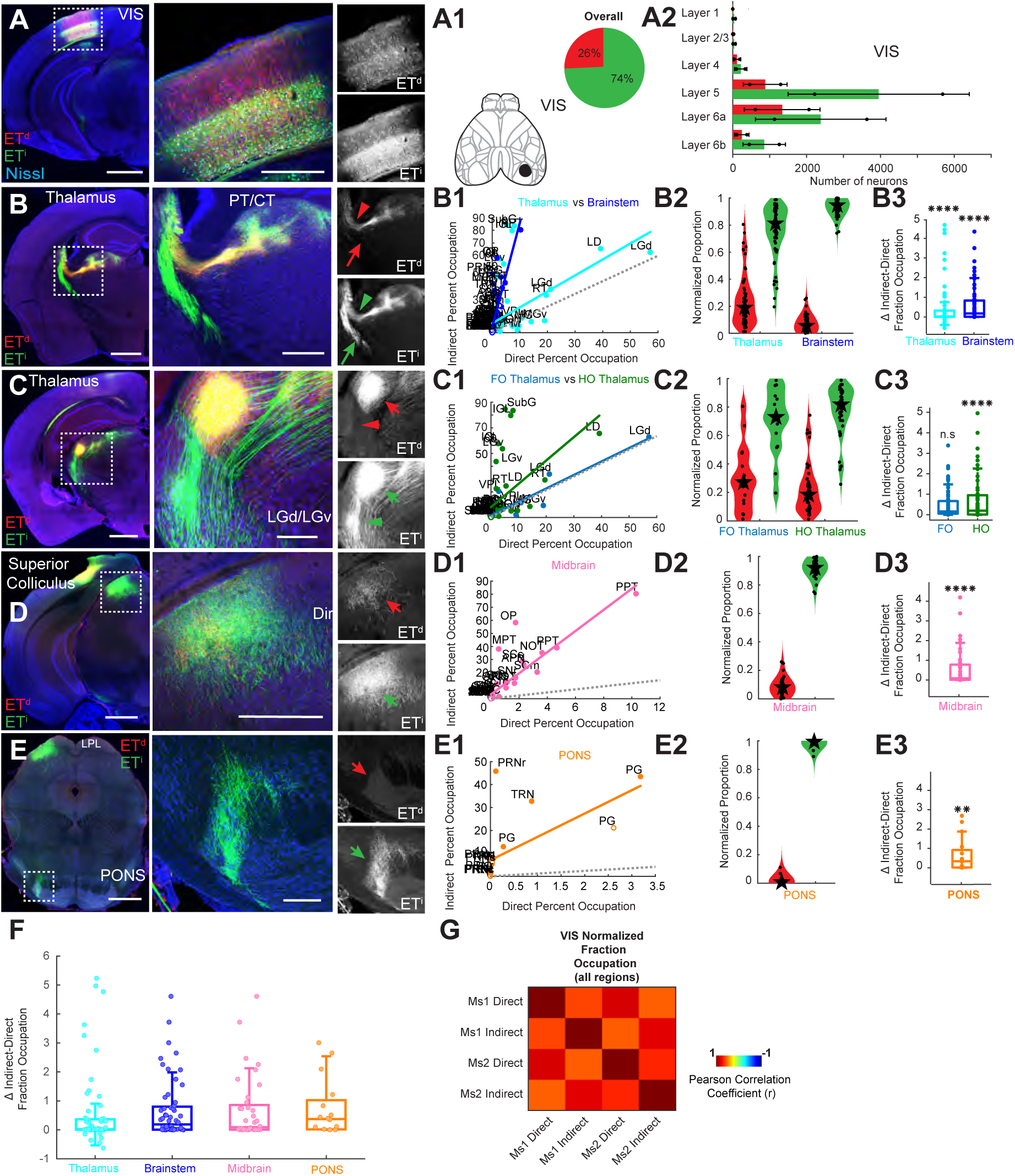
Predominant ET^i^ over ET^d^ subcortical projections from VIS, Related to Figure 4. (A) AAV injection in the VIS labeled ET^d^ (yellow) and ET^i^ (magenta). (A1) Left, location of the injection site shown on a schematic of dorsal cortex. Right, pie chart of proportion of ET^d^ and ET^i^ neurons labelled at the injection site. (A2) Bar plots of layer-wise counts of labelled ET^d^ and ET^i^ neurons in the injection site (n = 2 mice). (B) PT and CT fiber bundles with ET^d^ and ET^i^ projections from VIS. Notice both ET^d^ and ET^i^ projections in CT (arrowhead) while largely ET^i^ in PT (arrow). (C) Dense ET^d^ and ET^i^ projections to the thalamus, including LGd/LGv. (D) Stronger ET^i^ over ET^d^ projections to the superior colliculus. (E) Strong ET^i^ but nearly no ET^d^ projections to the PONS. (B1-E1) Scatter plots with linear regression of normalized fraction occupations of VIS ET^d^ and ET^i^ projections to target regions (B1) thalamus and brainstem, (C1) first order (FO) and higher order (HO) thalamus, (D1) midbrain, (E1) PONS. (B2-E2) Violin plots showing normalized percent occupation for VIS ET^d^ and ET^i^ projections with the star indicating the median and black circles showing individual values for subregions within target regions of (B2) thalamus and brainstem, (C2) first order (FO) and higher order (HO) thalamus, (D2) midbrain, (E2) PONS. (B3-E3) Box plots of Z-score normalized percent occupation differences between ET^i^ and ET^d^ projections from VIS to (B3) thalamus and brainstem, (C3) first order (FO) and higher order (HO) thalamus, (D3) midbrain, (E3) PONS. p-values reflect significant within region differences. (F) Box plots of Z-score normalized percent occupation differences between ET^i^ and ET^d^ projections from VIS for brainstem, midbrain and PONS with post-hoc comparisons to thalamus, following a multiple comparisons test (p < 0.01) to assess statistical differences among different projection target regions. There were no significant differences between target regions. (G) Correlation matrix heatmap showing pairwise Pearson correlations between ET^d^ and ET^i^ normalized fraction occupation across all areas for projections from VIS across two mice used for LightSheet Imaging. Abbreviations: VIS - visual cortex; FO – first order thalamic nuclei; HO- higher order thalamic nuclei; LGd - lateral geniculate dorsal; LGv – lateral geniculate ventral; PONS – pontine nuclei. Subregions for each target region are listed in Table S1. Scatter points in scatter plots, violin plots and box plots represent individual subregion percent occupation (normalized or raw) values, with filled markers for ipsilateral and open markers for contralateral data. Regression lines indicate overall correlation trends. Dotted grey line indicates the line of equality. Interquartile range in box plots is 25th–75th percentiles. Statistical tests used one-sample t-test, Wilcoxon signed-rank test, Kruskal–Wallis test, Wilcoxon rank-sum test and Kolmogorov-Smirnov test (p-values: *p < 0.05, **p < 0.01, ***p < 0.001, ****p < 0.0001, n.s = not significant) Scalebars in (A-E) 1 mm; (A-B right, D right) 500 µm; (C right) 250 µm; (E right) 200 µm.

**Figure S7.**
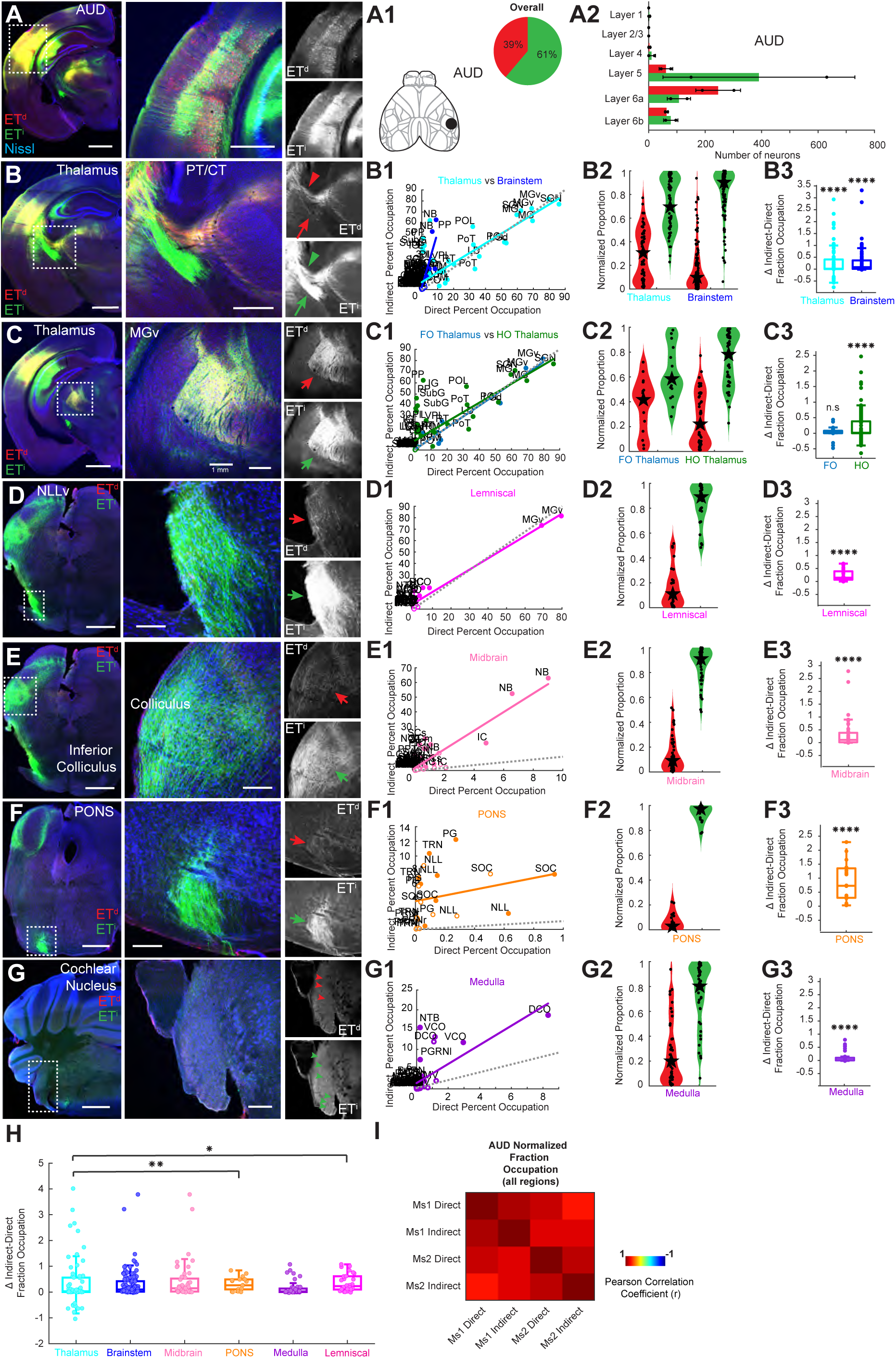
Predominant ET^i^ over ET^d^ subcortical projections from AUD, Related to Figure 4. (A) AAV injection in the AUD labeled ET^d^ (red) and ET^i^ (green). (A1) (left) Location of the injection site shown on a schematic of dorsal cortex. (right) Pie chart of proportion of ET^d^and ET^i^ neurons labelled at the injection site. (A2) Bar plots of layer-wise counts of labelled ET^d^ and ET^i^ neurons in the injection site (n = 2 mice). (B) PT and CT fiber bundles with ET^d^ and ET^i^ projections from AUD. Notice both ET^d^ and ET^i^ projections in CT (arrowhead) while largely ET^i^ in PT (arrow). (C) Dense ET^d^ and ET^i^ projections to the thalamus, including MGv. (D) Dense ET^i^ but weak ET^d^ projections to the NLLv. (E) Stronger ET^i^ over ET^d^ projections to the inferior colliculus. (F) Strong ET^i^ but weak ET^d^ projections to the PONS. (G) Sparse ET^i^ but nearly no ET^d^ projections to the CN. (B1-G1) Scatter plots with linear regression of normalized fraction occupations of AUD ET^d^ and ET^i^ projections to target regions (B1) thalamus and brainstem, (C1) first order (FO) and higher order (HO) thalamus, (D1) lemniscal sensory areas, (E1) midbrain, (F1) PONS, (G1) medulla. (B2-G2) Violin plots showing normalized percent occupation for AUD ET^d^ and ET^i^ projections with the Star indicating the median and black circles showing individual values for subregions within target regions of (B2) thalamus and brainstem (C2) first order (FO) and higher order (HO) thalamus (D2) lemniscal sensory areas (E2) midbrain (F2) PONS (G2) medulla. (B3-G3) Box plots of Z-score normalized percent occupation differences between ET^i^ and ET^d^ projections from AUD to (B3) thalamus and brainstem, (C3) first order (FO) and higher order (HO) thalamus, (D3) lemniscal sensory areas (E3) midbrain, (F3) PONS, (G3) medulla. p-values reflect significant within region differences. (H) Box plots of Z-score normalized percent occupation differences between ET^i^ and ET^d^ projections from AUD for brainstem, midbrain, PONS, medulla and lemniscal sensory areas with post-hoc comparisons to thalamus, following a multiple comparisons test (p < 0.01) to assess statistical differences among different projection target regions. The PONS and lemniscal sensory areas showed significantly more iNG-ET^i^ projections than thalamus. (I) Correlation matrix heatmap showing pairwise Pearson correlations between ET^d^ and ET^i^ normalized fraction occupation across all areas for projections from AUD across two mice used for LightSheet Imaging. Abbreviations: AUD – auditory cortex; FO – first order thalamic nuclei; HO- higher order thalamic nuclei; MGv – medial geniculate ventral; NLLv - nucleus of the lateral lemniscus ventral; PONS – pontine nuclei; BG – basal ganglia. Subregions for each target region are listed in Table S1. Scatter points in scatter plots, violin plots and box plots represent individual subregion percent occupation (normalized or raw) values, with filled markers for ipsilateral and open markers for contralateral data. Regression lines indicate overall correlation trends. Dotted grey line indicates the line of equality. Interquartile range in box plots is 25th–75th percentiles. Statistical tests used one-sample t-test, Wilcoxon signed-rank test, Kruskal–Wallis test, Wilcoxon rank-sum test and Kolmogorov-Smirnov test (p-values: *p < 0.05, **p < 0.01, ***p < 0.001, ****p < 0.0001, n.s = not significant) Scalebars in (A-G) 1 mm; (A-B right) 500 µm; (C right, E-G right) 200 µm; (D right) 100 µm.

**Figure S8.**
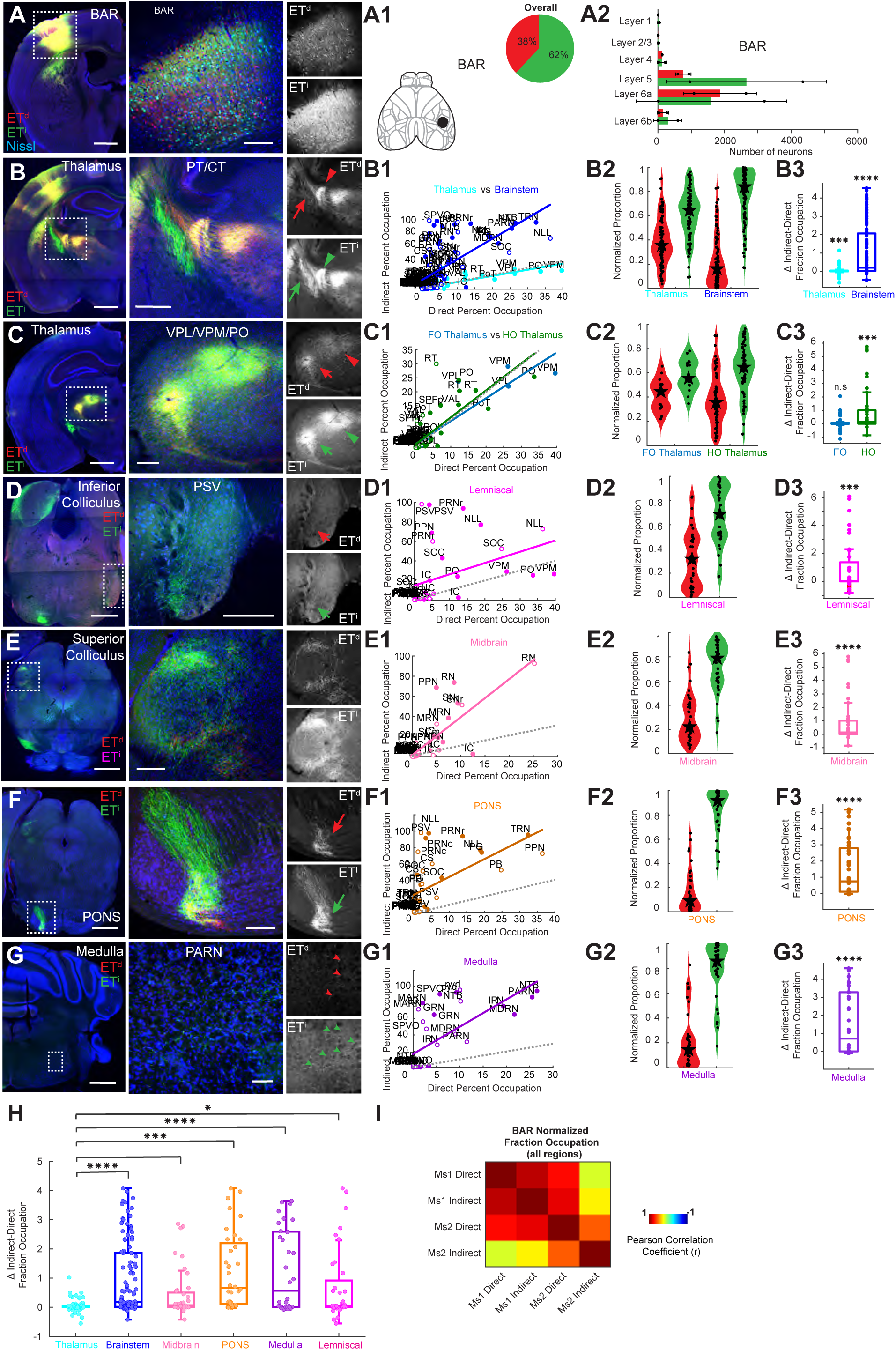
Predominant ET^i^ over ET^d^ subcortical projections from BAR, Related to Figure 4. (A) AAV injection in the BAR labeled ET^d^ (red) and ET^i^ (green). (A1) (left) Location of the injection site shown on a schematic of the dorsal cortex. (right) Pie chart of proportion of ET^d^and ET^i^ neurons labelled at the injection site. (A2) Bar plots of layer-wise counts of labelled ET^d^ and ET^i^ neurons in the injection site (n = 2 mice). (B) PT and CT fiber bundles with ET^d^ and ET^i^ projections from BAR. Notice both ET^d^ and ET^i^ projections in CT (arrowhead) while largely ET^i^ in PT (arrow). (C) Dense ET^d^ and ET^i^ projections to the thalamus, including VPL/VPM/PO. (D) Dense ET^i^ but weak ET^d^ projections to the inferior colliculus/PSV. (E) Stronger ET^i^ over ET^d^ projections to the superior colliculus. (F) Strong ET^i^ but weak ET^d^ projections to the PONS. (G) Strong ET^i^ but weak ET^d^ projections to the PARN. (B1-G1) Scatter plots with linear regression of normalized fraction occupations of BAR ET^d^ and ET^i^ projections to target regions (B1) thalamus and brainstem, (C1) first order (FO) and higher order (HO) thalamus, (D1) lemniscal sensory areas, (E1) midbrain, (F1) PONS, (G1) medulla. (B2-G2) Violin plots showing normalized percent occupation for BAR ET^d^ and ET^i^ projections with the Star indicating the median and black circles showing individual values for subregions within target regions of (B2) thalamus and brainstem (C2) first order (FO) and higher order (HO) thalamus (D2) lemniscal sensory areas (E2) midbrain (F2) PONS (G2) medulla. (B3-G3) Box plots of Z-score normalized percent occupation differences between ET^i^ and ET^d^ projections from BAR to (B3) thalamus and brainstem, (C3) first order (FO) and higher order (HO) thalamus, (D3) lemniscal sensory areas (E3) midbrain, (F3) PONS, (G3) medulla. p-values reflect significant within region differences. (H) Box plots of Z-score normalized percent occupation differences between ET^i^ and ET^d^ projections from BAR for brainstem, midbrain, PONS, medulla and lemniscal sensory areas with post-hoc comparisons to thalamus, following a multiple comparisons test (p < 0.0001) to assess statistical differences among different projection target regions. The brainstem, midbrain, PONS, medulla and lemniscal sensory areas showed significantly more iNG-ET^i^ projections than thalamus. (I) Correlation matrix heatmap showing pairwise Pearson correlations between ET^d^ and ET^i^ normalized fraction occupation across all areas for projections from BAR across two mice used for LightSheet Imaging. Abbreviations: BAR – barrel cortex; FO – first order thalamic nuclei; HO- higher order thalamic nuclei; VPM – ventral posterior medial nucleus; VPL – ventral posterior lateral nucleus; PO – posterior thalamic nucleus; PONS – pontine nuclei; PSV – principal sensory nucleus of the trigeminal. Subregions for each target region are listed in Table S1. Scatter points in scatter plots, violin plots and box plots represent individual subregion percent occupation (normalized or raw) values, with filled markers for ipsilateral and open markers for contralateral data. Regression lines indicate overall correlation trends. Dotted grey line indicates the line of equality. Interquartile range in box plots is 25th–75th percentiles. Statistical tests used one-sample t-test, Wilcoxon signed-rank test, Kruskal–Wallis test, Wilcoxon rank-sum test and Kolmogorov-Smirnov test (p-values: *p < 0.05, **p < 0.01, ***p < 0.001, ****p < 0.0001, n.s = not significant) Scalebars in (A-G) 1 mm; (B right) 500 µm; (A right, C-F right) 200 µm; (G right) 100 µm.

**Figure S9.**
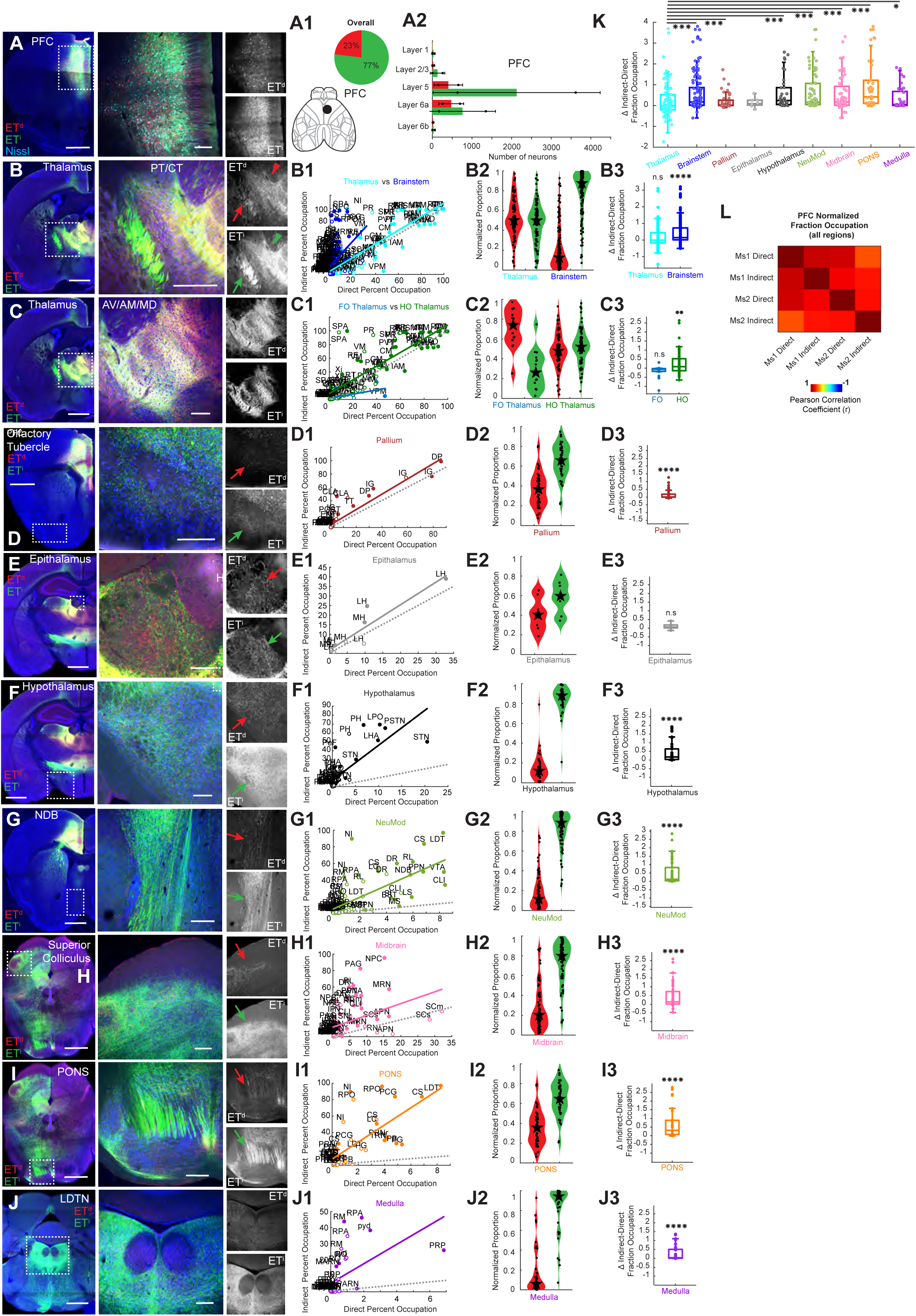
Predominant ET^i^ over ET^d^ subcortical projections from PFC, Related to Figure 5. (A) AAV injection in the PFC labeled ET^d^ (red) and ET^i^ (green). (A1) (left) Location of the injection site shown on a schematic of dorsal cortex. (right) Pie chart of proportion of ET^d^and ET^i^ neurons labelled at the injection site. (A2) Bar plots of layer-wise counts of labelled ET^d^ and ET^i^ neurons in the injection site (n = 2 mice). (B) PT and CT fiber bundles with ET^d^ and ET^i^ projections from PFC. Notice both ET^d^ and ET^i^ projections in CT (arrowhead) while largely ET^i^ in PT (arrow). (C) Dense ET^d^ and ET^i^ projections to the thalamus, including AV/AL/MD. (D) Dense ET^i^ but weak ET^d^ projections to the olfactory tubercle. (E) Similar ET^i^ and ET^d^ projections to the epithalamus. (F) Strong ET^i^ but weak ET^d^ projections to the hypothalamus. (G) Strong ET^i^ but weak ET^d^ projections to the NDB. (H) Strong ET^i^ but weak ET^d^ projections to the superior colliculus. (I) Strong ET^i^ but weak ET^d^ projections to the PONS. (J) Strong ET^i^ but nearly no ET^d^ projections to the LDTN. (B1-J1) Scatter plots with linear regression of normalized fraction occupations of PFC ET^d^ and ET^i^ projections to target regions (B1) thalamus and brainstem, (C1) first order (FO) and higher order (HO) thalamus, (D1) pallial areas, (E1) epithalamus, (F1) hypothalamus, (G1) neuromodulatory structures, (H1) midbrain, (I1) PONS, (J1) medulla. (B2-J2) Violin plots showing normalized percent occupation for PFC ET^d^ and ET^i^ projections with the Star indicating the median and black circles showing individual values for subregions within target regions of (B2) thalamus and brainstem, (C2) first order (FO) and higher order (HO) thalamus, (D2) pallial areas, (E2) epithalamus, (F2) hypothalamus, (G2) neuromodulatory structures, (H2) midbrain, (I2) PONS, (J2) medulla. (B3-J3) Box plots of Z-score normalized percent occupation differences between ET^i^ and ET^d^ projections from PFC to (B3) thalamus and brainstem, (C3) first order (FO) and higher order (HO) thalamus, (D3) pallial areas, (E3) epithalamus, (F3) hypothalamus, (G3) neuromodulatory structures, (H3) midbrain, (I2) PONS, (J3) medulla. p-values reflect significant within region differences. (K) Box plots of Z-score normalized percent occupation differences between ET^i^ and ET^d^ projections from PFC for brainstem, pallium, epithalamus, hypothalamus, neuromodulatory structures, midbrain, PONS and medulla with post-hoc comparisons to thalamus, following a multiple comparisons test (p < 0.0001) to assess statistical differences among different projection target regions. The brainstem, pallium, hypothalamus, neuromodulatory structures, midbrain, PONS and medulla areas showed significantly more iNG-ET^i^ projections than thalamus. (L) Correlation matrix heatmap showing pairwise Pearson correlations between ET^d^ and ET^i^ normalized fraction occupation across all areas for projections from PFC across two mice used for LightSheet Imaging. Abbreviations: PFC – prefrontal cortex; FO – first order thalamic nuclei; HO- higher order thalamic nuclei; VPM – ventral posterior medial nucleus; AV – anteroventral nucleus; AM – anteromedial nucleus; MD – mediodorsal nucleus of thalamus; NDB – nucleus of the diagonal band; LDTN – lateral dorsal tegmental nucleus; PONS – pontine nuclei. Subregions for each target region are listed in Table S1. Scatter points in scatter plots, violin plots and box plots represent individual subregion percent occupation (normalized or raw) values, with filled markers for ipsilateral and open markers for contralateral data. Regression lines indicate overall correlation trends. Dotted grey line indicates the line of equality. Interquartile range in box plots is 25th–75th percentiles. Statistical tests used one-sample t-test, Wilcoxon signed-rank test, Kruskal–Wallis test, Wilcoxon rank-sum test and Kolmogorov-Smirnov test (p-values: *p < 0.05, **p < 0.01, ***p < 0.001, ****p < 0.0001, n.s = not significant) Scalebars in (A-J) 1 mm; (B right) 500 µm; (A right, C-D right, F-J right) 200 µm; (E right) 100 µm.

**Figure S10.**
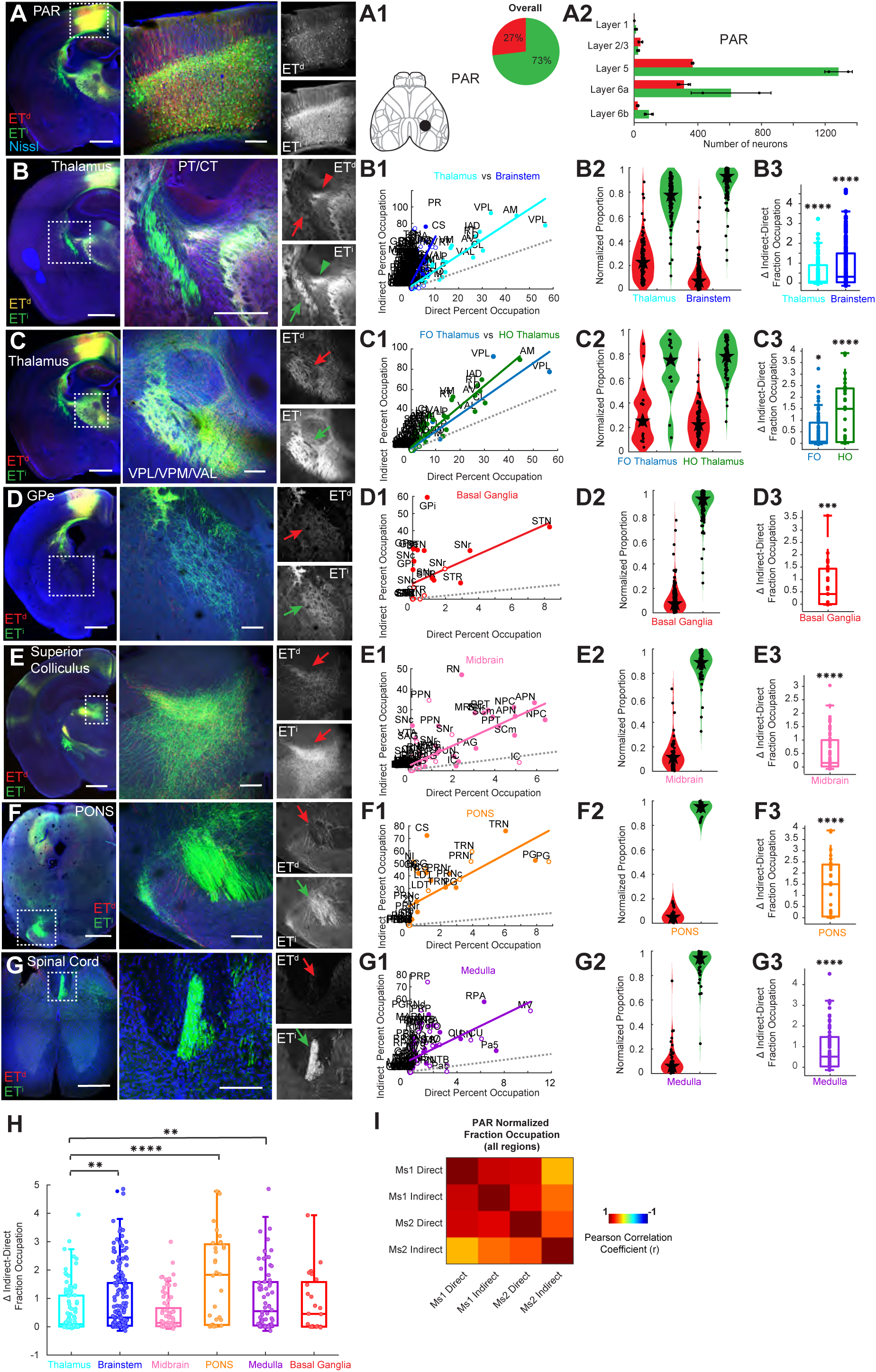
Predominant ET^i^ over ET^d^ subcortical projections from PAR, Related to Figure 5. (A) AAV injection in the PAR labeled ET^d^ (red) and ET^i^ (green). (A1) (left) Location of the injection site shown on a schematic of dorsal cortex. (right) Pie chart of proportion of ET^d^and ET^i^ neurons labelled at the injection site. (A2) Bar plots of layer-wise counts of labelled ET^d^ and ET^i^ neurons in the injection site (n = 2 mice). (B) PT and CT fiber bundles with ET^d^ and ET^i^ projections from PAR. Notice both ET^d^ and ET^i^ projections in CT (arrowhead) while largely ET^i^ in PT (arrow). (C) Dense ET^d^ and ET^i^ projections to the thalamus, including VPL/VPM/VA. (D) Dense ET^i^ but weak ET^d^ projections to the GPe. (E) Stronger ET^i^ over ET^d^ projections to the superior colliculus. (F) Strong ET^i^ but weak ET^d^ projections to the PONS. (G) Strong ET^i^ but nearly no ET^d^ projections to the spinal cord. (B1-G1) Scatter plots with linear regression of normalized fraction occupations of PAR ET^d^ and ET^i^ projections to target regions (B1) thalamus and brainstem, (C1) first order (FO) and higher order (HO) thalamus, (D1) basal ganglia, (E1) midbrain, (F1) PONS, (G1) medulla. (B2-G2) Violin plots showing normalized percent occupation for PAR ET^d^ and ET^i^ projections with the Star indicating the median and black circles showing individual values for subregions within target regions of (B2) thalamus and brainstem (C2) first order (FO) and higher order (HO) thalamus (D2) basal ganglia (E2) midbrain (F2) PONS (G2) medulla. (B3-G3) Box plots of Z-score normalized percent occupation differences between ET^i^ and ET^d^ projections from PAR to (B3) thalamus and brainstem, (C3) first order (FO) and higher order (HO) thalamus, (D3) basal ganglia, (E3) midbrain, (F3) PONS, (G3) medulla. p-values reflect significant within region differences. (H) Box plots of Z-score normalized percent occupation differences between ET^i^ and ET^d^ projections from PAR for brainstem, midbrain, PONS, medulla and BG with post-hoc comparisons to thalamus, following a multiple comparisons test (p < 0.001) to assess statistical differences among different projection target regions. The brainstem, PONS and medulla regions showed significantly more iNG-ET^i^ projections than thalamus. (I) Correlation matrix heatmap showing pairwise Pearson correlations between ET^d^ and ET^i^ normalized fraction occupation across all areas for projections from PAR across two mice used for LightSheet Imaging. Abbreviations: PAR – parietal cortex; FO – first order thalamic nuclei; HO- higher order thalamic nuclei; VPM – ventral posterior medial nucleus; VPL – ventral posterior lateral nucleus; VAL – ventral anterio-lateral complex of thalamus; GPe – globus pallidus externa; PONS – pontine nuclei; BG – basal ganglia. Subregions for each target region are listed in Table S1. Scatter points in scatter plots, violin plots and box plots represent individual subregion percent occupation (normalized or raw) values, with filled markers for ipsilateral and open markers for contralateral data. Regression lines indicate overall correlation trends. Dotted grey line indicates the line of equality. Interquartile range in box plots is 25th–75th percentiles. Statistical tests used one-sample t-test, Wilcoxon signed-rank test, Kruskal–Wallis test, Wilcoxon rank-sum test and Kolmogorov-Smirnov test (p-values: *p < 0.05, **p < 0.01, ***p < 0.001, ****p < 0.0001, n.s = not significant) Scalebars in (A-G) 1 mm; (B right) 500 µm; (G right) 400 µm; (A right,C-F right) 200 µm.

**Figure S11.**
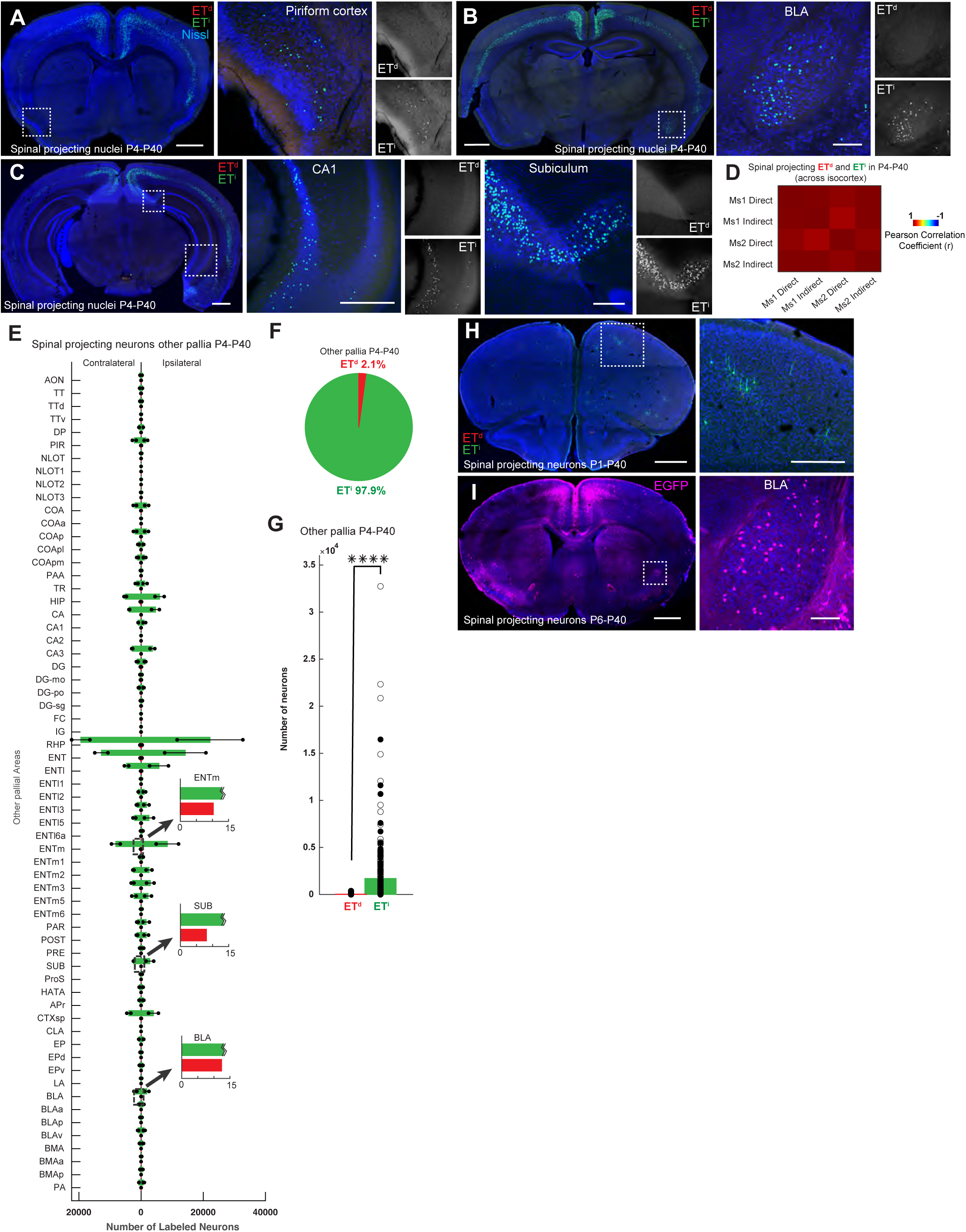
Developing ET^i^ neurons across cerebral cortical structures initially all project axons to the spinal cord. Related to Figure 6. (A-C) Spinal-projecting neurons retrogradely labelled from injections to cSpC of P4 mice and harvested at P40 (P4-P40) in (A) piriform cortex (B) BLA (C) Hippocampus – CA1 & subiculum. Note almost all of the labelled nuclei are ET^i^. (D) Correlation matrix heatmap showing pairwise Pearson correlations between ET^d^ and ET^i^ nuclei retrogradely labelled from cSpC in P4-P40 mice, across different non-isocortical pallial areas from two mice used for LightSheet Imaging. (E) Bar plots of retrogradely labelled spinal-projecting ET^d^ (red) and ET^i^ (green) nuclei numbers across different non-isocortical pallial areas labelled from injections into cSpC in P4-P40 mice. Zoomed in sections (dotted black box) of the bar plots showing small to zero number of ET^d^ with the longer bar of ET^i^ which exceeds the zoomed scale truncated. (F) Proportions of spinal-projecting ET^d^ (red) and ET^i^ (green) nuclei across different non-isocortical pallial areas in P4-P40 mice. (G) Number of retrogradely labelled corticospinal ET^i^ are significantly more than ET^d^ across different non-isocortical pallial areas in P4-P40 mice. (H) Sparse labelling of retrogradely labelled spinal-projecting ET^d^ and ET^i^ neurons in the isocortex from injections into cSpC in P1-P40 mice. (I) Retrogradely labelled spinal-projecting ET^d^ and ET^i^ neurons in the isocortex and BLA from injections of generic retro-AAV into cSpC of P6 wildtype mice and harvested at P40 (P6-P40). Abbreviations: BLA – basolateral amygdala; CA1 – Cornu Ammonis area 1; cSpC – cervical spinal cord. P1-P40 – injected in P1 and harvested at P40; P4-P40 – injected in P4 and harvested at P40; P28-P40 – injected in P28 and harvested at P40 Subregions in (E) are listed in Table S1. Statistical tests used: two-sample t-test, (p-values: ****p < 0.0001) Scalebars in (A-C, H-I) 1 mm; (C middle, H right) 500 µm; (B right, C right, I right) 200 µm.

## Supplementary Videos

**Supplementary Video 1.** Whole-brain Lightsheet image stack of retrogradely labelled ET^d^ (red) and ET^i^ (green) neurons across the isocortex from injections into the superior colliculus(SC) in a FezF2TC;Tbr2Flp mouse.

**Supplementary Video 2.** Whole-brain Lightsheet image stack of retrogradely labelled ET^d^ (red) and ET^i^ (green) neurons across the isocortex from injections into the pontine nuclei (PONS) in a FezF2TC;Tbr2Flp mouse.

**Supplementary Video 3.** Whole-brain Lightsheet image stack of retrogradely labelled ET^d^ (red) and ET^i^ (green) neurons across the isocortex from injections into the medulla (MED) in a FezF2TC;Tbr2Flp.

**Supplementary Video 4.** Whole-brain Lightsheet image stack of retrogradely labelled ET^d^ (red) and ET^i^ (green) neurons across the isocortex from injections into the cervical spinal cord (cSpC) in a FezF2TC;Tbr2Flp mouse.

**Supplementary Video 5.** Whole-brain Lightsheet image stack of anterograde axonal projections of ET^d^ (red) and ET^i^ (green) neurons in the rostral forelimb orofacial area (RFO) of the isocortex in a FezF2TC;Tbr2Flp mouse.

**Supplementary Video 6.** Whole-brain Lightsheet image stack of anterograde axonal projections of ET^d^ (red) and ET^i^ (green) neurons in the caudal forelimb area (CFA) of the isocortex in a FezF2TC;Tbr2Flp mouse.

**Supplementary Video 7.** Whole-brain Lightsheet image stack of anterograde axonal projections of ET^d^ (red) and ET^i^ (green) neurons in the secondary motor area (MOs) of the isocortex in a FezF2TC;Tbr2Flp mouse.

**Supplementary Video 8.** Whole-brain Lightsheet image stack of anterograde axonal projections of ET^d^ (red) and ET^i^ (green) neurons in the visual cortex (VIS) in a FezF2TC;Tbr2Flp mouse.

**Supplementary Video 9.** Whole-brain Lightsheet image stack of anterograde axonal projections of ET^d^ (red) and ET^i^ (green) neurons in the auditory cortex (AUD) in a FezF2TC;Tbr2Flp mouse.

**Supplementary Video 10.** Whole-brain Lightsheet image stack of anterograde axonal projections of ET^d^ (red) and ET^i^ (green) neurons in the barrel cortex (BAR) in a FezF2TC;Tbr2Flp mouse.

**Supplementary Video 11.** Whole-brain Lightsheet image stack of anterograde axonal projections of ET^d^ (red) and ET^i^ (green) neurons in the prefrontal cortex (PFC) in a FezF2TC;Tbr2Flp mouse.

**Supplementary Video 12.** Whole-brain Lightsheet image stack of anterograde axonal projections of ET^d^ (red) and ET^i^ (green) neurons in the parietal cortex (PAR) in a FezF2TC;Tbr2Flp mouse.

**Supplementary Video 13.** Whole-brain Lightsheet image stack of retrogradely labelled ET^d^ (red) and ET^i^(green) nuclei across different cortical/pallial areas from injections into the cervical spinal cord (cSpC) of a P4 FezF2TC;Tbr2Flp mouse and harvested at P40 (P4-P40).

**Supplementary Video 14.** Behavioural effects of optogenetic activation of ET^all^,ET^d^ and ET^i^ in the RFO shown sequentially in both the head-fixed and free moving conditions.

## Notes

https://github.com/ShreyasDuke/ETd-ETi-Code.git

### Competing Interest Statement

The authors have declared no competing interest.

### Summary of Updates

Manuscript revised with new data. Figures and text updated.

